# Comprehensive preclinical evaluation of human-derived anti-poly-GA antibodies in cellular and animal models of C9ORF72 disease

**DOI:** 10.1101/2022.01.13.475329

**Authors:** Melanie Jambeau, Kevin D. Meyer, Marian Hruska-Plochan, Ricardos Tabet, Chao-Zong Lee, Ananya Ray-Soni, Corey Aguilar, Kitty Savage, Nibha Mishra, Nicole Cavegn, Petra Borter, Chun-Chia Lin, Karen Jansen-West, Jay Jiang, Fernande Freyermuth, Nan Li, Pierre De Rossi, Manuela Pérez-Berlanga, Xin Jiang, Lilian M. Daughrity, Joao Pereira, Sarav Narayanan, Yuanzheng Gu, Shekhar Dhokai, Isin Dalkilic-Liddle, Zuzanna Maniecka, Julien Weber, Michael Workman, Melissa McAlonis-Downes, Eugene Berezovski, Yong-Jie Zhang, James Berry, Brian J. Wainger, Mark W. Kankel, Mia Rushe, Christoph Hock, Roger M. Nitsch, Don W. Cleveland, Leonard Petrucelli, Tania Gendron, Fabio Montrasio, Jan Grimm, Magdalini Polymenidou, Clotilde Lagier-Tourenne

**Affiliations:** Department of Neurology, The Sean M. Healey and AMG Center for ALS at Mass General, Massachusetts General Hospital and Harvard Medical School, Boston, MA 02114, USA; Broad Institute of Harvard and MIT, Cambridge, MA 02142, USA; Department of Quantitative Biomedicine, University of Zurich, Winterthurerstrasse 190, CH-8057 Zurich, Switzerland; Neurimmune AG, Wagistrasse 18, CH-8952 Schlieren, Switzerland; Department of Neuroscience, Mayo Clinic, Jacksonville, FL 32224, USA; Ludwig Institute for Cancer Research, University of California San Diego, La Jolla, CA 92093; Biogen, Cambridge, Massachusetts, 02142, USA; Institute for Regenerative Medicine, University of Zurich, Wagistrasse 12, CH-8952 Schlieren, Switzerland

## Abstract

Hexanucleotide G_4_C_2_ repeat expansions in the *C9ORF72* gene are the most common genetic cause of amyotrophic lateral sclerosis (ALS) and frontotemporal dementia (FTD). Dipeptide repeat proteins (DPRs) generated by translation of repeat-containing RNAs show toxic effects *in vivo* as well as *in vitro* and are key targets for therapeutic intervention. We generated human antibodies that bind DPRs with high affinity and specificity. Anti-GA antibodies engaged extra- and intracellular poly-GA and reduced aggregate formation in a poly-GA over-expressing human cell line. However, antibody treatment in human neuronal cultures synthesizing exogenous poly-GA resulted in the formation of large extracellular immune complexes and did not affect accumulation of intracellular poly-GA aggregates. Treatment with antibodies was also shown to directly alter the morphological and biochemical properties of poly-GA and to shift poly-GA/antibody complexes to more rapidly sedimenting ones. These alterations were not observed with poly-GP and have important implications for accurate measurement of poly-GA levels including the need to evaluate all centrifugation fractions and disrupt the interaction between treatment antibodies and poly-GA by denaturation. Targeting poly-GA and poly-GP in two mouse models expressing G_4_C_2_ repeats by systemic antibody delivery for up to 16 months was well-tolerated and led to measurable brain penetration of antibodies. Long term treatment with anti-GA antibodies produced improvement in an open field movement test in aged C9ORF72^450^ mice. However, chronic administration of anti-GA antibodies in AAV-(G_4_C_2_)_149_ mice was associated with increased levels of poly-GA detected by immunoassay and did not significantly reduce poly-GA aggregates or alleviate disease progression in this model.

**Significance:** Immunotherapy has been proposed for neurodegenerative disorders including Alzheimer’s or Parkinson’s diseases. Recent reports using antibodies against poly-GA or active immunization suggested similar immunotherapy in ALS/FTD caused by repeat expansion in the C9ORF72 gene (1, 2). Here, we systematically characterized human antibodies against multiple DPR species and tested the biological effects of antibodies targeting poly-GA in different cellular and mouse models. Target engagement was shown in three independent cellular models. Anti-GA antibodies reduced the number of intracellular poly-GA aggregates in human T98G cells but not in cultured human neurons. Whereas chronic anti-GA treatment in BAC C9ORF72^450^ mice did not impact poly-GA levels and modestly improved one behavioral phenotype, poly-GA levels detected by immunoassays were increased and disease progression was unaltered in AAV-(G_4_C_2_)_149_ mice.

## Introduction

Hexanucleotide repeat expansions (G_4_C_2_) in the *C9ORF72* gene are the most frequent genetic cause of amyotrophic lateral sclerosis (ALS) and frontotemporal dementia (FTD) (3, 4). Proposed disease mechanisms include *C9ORF72* haploinsufficiency, repeat-RNA toxicity and protein toxicity. Though the relative contribution of each mechanism is not fully understood (5), there is mounting evidence that accumulation of dipeptide repeat proteins (DPRs), generated by repeat-associated non-ATG (RAN) translation across the *C9ORF72* expansion, plays a crucial role in neurodegeneration (6–11). DPRs are translated from both sense (poly-GA, poly-GR and poly-GP) and antisense (poly-PR, poly-PA and poly-GP) repeat-containing RNAs, and represent the major component of p62-positive, TDP-43-negative aggregates in the central nervous system of C9ORF72 ALS/FTD patients (12–14). Moreover, several *in vivo* and *in vitro* studies support direct toxic effects of arginine-rich DPR proteins poly-PR and poly-GR (15–19), as well as the aggregation-prone and most abundant DPR product, poly-GA (20–22). A recent study directly comparing congenic mice expressing either poly-GA or poly-PR indicates that poly-GA is considerably more toxic, leading to TDP-43 abnormalities and neuronal loss (23), highlighting its suitability as a therapeutic target.

Passive immunotherapy using humanized or fully human antibodies targeting aberrantly produced or misfolded proteins has been investigated in several pre-clinical and clinical settings for the treatment of neurodegenerative disorders (24, 25). The most advanced programs have targeted extracellular amyloid-β plaques in mouse models and patients with Alzheimer’s disease (26, 27). Targeting intracellular misfolded proteins, including tau and α-synuclein, has also shown beneficial effects on pathology and behavioral abnormalities in multiple mouse models of Alzheimer’s (28, 29) or Parkinson’s diseases (30). Recently, human-derived antibodies targeting misfolded SOD1 were reported to delay disease onset and increase survival in independent ALS-linked SOD1 mutant mouse models (31). Another study explored the potential of immunotherapy for treating C9ORF72 ALS/FTD by using a mouse-derived antibody against poly-GA in cultured cells (32), and a potential beneficial effect of poly-GA human-derived antibodies was reported in C9ORF72 BAC transgenic mice expressing 500 repeats (1). Most recently, the effect of active immunization against poly-GA (2) was assessed in a mouse model overexpressing poly-GA fused with the cyan fluorescent protein ((GA)_149_-CFP;). Reduction in poly-GA accumulation, neuroinflammation and TDP-43 mislocalization was observed in (GA)_149_-CFP mice immunized with ovalbumin-(GA)10 conjugates (2). Several non-exclusive mechanisms have been proposed for antibody-mediated neutralization of intra-cellular aggregates. In particular, antibodies to tau or α-synuclein may influence disease progression by inhibiting cell-to-cell propagation of toxic proteins (29, 33), a mechanism proposed also for C9ORF72 DPRs (22, 32, 34). Moreover, several studies suggest that antibodies are internalized by neuronal cells (35, 36), where they may capture accumulated protein aggregates and facilitate their degradation.

Here, we systematically characterized 11 human anti-DPR antibodies generated by immune repertoire analyses of healthy elderly donors, and tested poly-GA antibodies in multiple cell lines, including human neuronal cultures and two C9ORF72 mouse models. Poly-GA-specific antibodies entered cultured neurons and colocalized with their target in intracellular vesicles. Moreover, long-term antibody treatment of human neurons expressing poly-GA resulted in capturing of extracellular poly-GA and lead to the formation of extracellular antibody-poly-GA complexes. In transgenic mice expressing the *C9ORF72* gene containing 450 G_4_C_2_ repeats (C9^450^) or in mice expressing 149 G_4_C_2_ repeats within the central nervous system by means of adeno-associated virus (AAV-G_4_C_2_) (37, 38), antibodies were shown to cross the blood-brain barrier without obvious adverse effects upon long term chronic administration. However, in the (AAV-G_4_C_2_)_149_ mice antibody treatment was not efficient in clearing poly-GA aggregates and was associated with increased poly-GA levels measured by immunoassay. While an improvement was observed in one behavioral assay in C9^450^ mice, treatment was not associated with alleviation of disease progression in (AAV-G_4_C_2_)_149_ mice.

## Results

### Antibody generation and affinity determination

Human monoclonal antibodies targeting the five *C9ORF72* DPRs were generated by screening memory B-cell libraries from healthy elderly subjects, an approach previously used to identify potent antibodies recognizing protein aggregates that include amyloid-β, SOD1 or α-synuclein (25). Eleven antibodies with high affinity to one or multiple DPRs were characterized by ELISA *(SI Appendix,* **Table S1**, **Fig. SL*A***), biolayer interferometry *(SI Appendix,* **Table S1**, **Fig. S1*B***) and immunostaining (*SI Appendix,* **Table S1**, **Fig. S2-6**). Four poly-GA-specific antibodies, designated α-GA_1-4_, were identified with nanomolar EC_50_ constants (0.2-0.3 nM) (*SI Appendix,* **Table S1**, **Fig. S1*A***). Kinetic analyses by biolayer interferometry revealed that the four α-GA antibodies had comparable association rate constants (*k*_a_) to GA_15_ peptides. α-GA_2-4_ showed comparably low dissociation rates (*k*_d_), whereas faster target dissociation was observed for α-GA_1_ (*SI Appendix,* **Table S1**, **Fig. S1*B***). Antibody α-GP_1_ displayed high affinity binding to poly-GP and a 26-fold lower affinity to poly-GA (*SI Appendix,* **Table S1**, **Fig. S1*A***). By screening against the arginine-rich DPR proteins, poly-GR and poly-PR, we further identified α-PR_2_, which exclusively recognized poly-PR (EC_50_ of 12.8 nM), as well as several antibody candidates targeting more than one DPR species, α-PR_1,3_ and α-GR_1_. Two candidates with high affinity EC_50_ binding (0.1-0.4 nM) to poly-PA were identified, with α-PA_1_ specifically targeting poly-PA (*SI Appendix,* **Table S1, Fig. S1*A***).

The four human anti-GA (α-GA_1-4_) and the anti-GP (α-GP_1_) antibodies specifically recognized aggregates in human brain tissues from ALS patients carrying pathogenic expansions in the *C9ORF72* gene *(SI Appendix,* **Table S1**, **Fig. S2*A*** and ***B***, upper panels) and in transgenic mice expressing 450 *C9ORF72* hexanucleotide repeats (C9^450^) (37) (*SI Appendix,* **Table S1, Fig. S2*A*** and ***B***, lower panels). Antibodies targeting poly-GR, poly-PR and poly-PA (α-GR_1_, α-PR_1-3_ and α-PA_1,2_) failed to detect DPR aggregates following immunostaining of formalin-fixed human and mouse tissues (*SI Appendix,* **Table S1**).

To further test antibody specificity across all five DPRs, we transiently transfected motor neuron-like cells (NSC-34) with single DPR species with 50 repeats and tagged with the enhanced green fluorescent protein (GFP) (*SI Appendix,* **Fig. S2-6**). Immunofluorescence analysis revealed a predominantly cytoplasmic, diffuse distribution of GA_50_-GFP, GP_47_-GFP and PA_5_0-GFP, with GA_50_-GFP also forming dense and bright aggregates. GR_50_-GFP accumulated either in the cytoplasm or in the nucleus and PR_50_-GFP localized in nuclei. All four human-derived α-GA antibodies (α-GA_1-4_) specifically recognized poly-GA, with no cross-reactivity to other DPR species (*SI Appendix,* **Table S1, Fig. S2 *C*** and ***D*, Fig. S3**). Specificity of the α-GA antibodies was confirmed by absence of signal in GFP-only transfected cells and in cells stained with secondary antibody only (*SI Appendix,* **Fig. S4*A***). In addition to displaying a strong, specific staining for poly-GP, antibody α-GP_1_ also showed a weak reactivity for poly-GA, consistent with its *in vitro* binding affinity (EC_50_ values in *SI Appendix,* **Table S1**), as well as a faint non-specific nuclear staining (*SI Appendix,* **Fig. S4*B***). Two α-PA antibodies, α-PA_1,2_, specifically recognized poly-PA (*SI Appendix,* **Table S1, Fig. S5**), while antibodies α-PR_2,3_ detected poly-PR in the nucleus, particularly in nucleoli, but no other DPR species (*SI Appendix,* **Table S1, Fig. S6**). Since α-GA and α-GP reliably detected aggregates in C9^450^ mouse and *C9ORF72* ALS patient brain sections, we selected α-GA_1_, α-GA_3_ and α-GP_1_ (or murine chimeric IgG2a derivatives of each: ^ch^α-GA_1_, ^ch^α-GA_3_ and ^ch^α-GP1) to test their ability to impact poly-GA and poly-GP accumulation *in vitro* and *in vivo*.

### Antibody uptake and colocalization with poly-GA in living cells

To determine if living cells internalize anti-GA antibodies, human neuroblastoma SH-SY5Y cells were transfected to express either a GFP control construct or GA_50_-GFP and incubated in media containing antibodies α-GA_1_ α-GA_3_ or an IgG isotype control (50 nM, 72 hrs). A strong antibody signal was detected in SH-SY5Y cells expressing GA_50_-GFP and treated with α-GA_1_ or α-GA_3_ compared to cells incubated with an IgG isotype control or cells expressing GFP only (**Fig. *1A*** and ***B**, SI Appendix*, **Fig. S7 *A*** and ***B***). This result is consistent with the presence of poly-GA within cells enhancing retention of internalized α-GA human antibodies. Automated quantification confirmed colocalization between GA_50_-GFP and α-GA antibodies. Indeed, 43 % and 55 % of GA_50_-GFP area colocalized with α-GA_1_ and α-GA_3_, respectively, while less than 2 % of the GFP-positive area colocalized with these antibodies (p < 0.001; **Fig. 1*B***). Importantly, GA_50_-GFP did not colocalize with the IgG isotype control (p < 0.001; **Fig. 1*B***). Similar results were obtained when quantifying the total area of antibodies that colocalized with GFP versus GA_50_-GFP, with approximately 40 % of the α-GA_1_ and α-GA_3_ signal overlapping with GA_50_-GFP, while less than 8 % overlapped with GFP (p < 0.001; *SI Appendix,* **Fig. S7*B***).

**Fig. 1.**
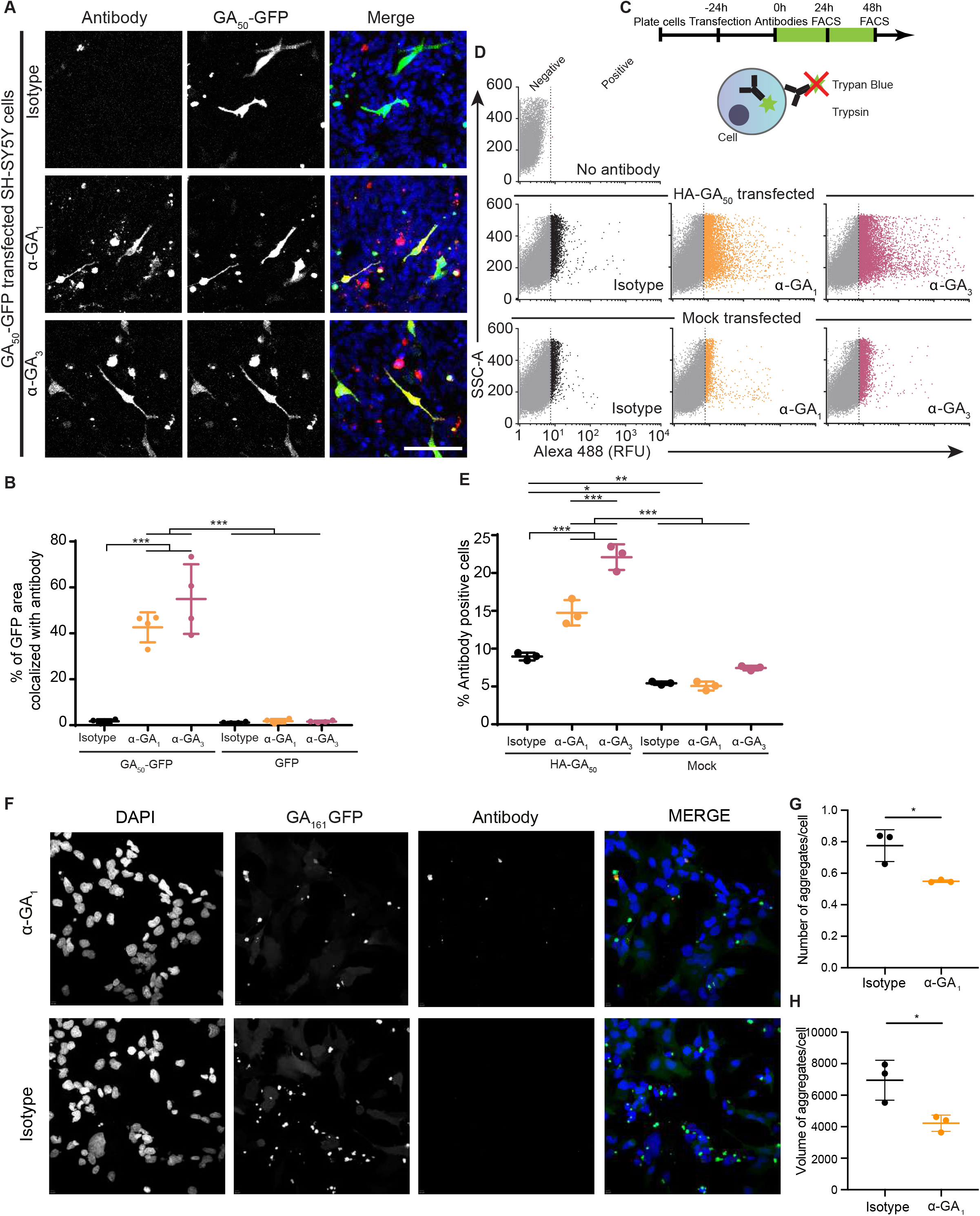
Antibody uptake and poly-GA colocalization in living cells. **(A)** Confocal fluorescence images of SH-SY5Y cells transfected with GA_50_-GFP (green) and incubated with human α-GA_1_. α-GA_3_ or an IgG control antibody (72 hrs). Antibodies were visualized after fixation with a secondary α-human IgG antibody (red) and nuclei with DAPI (blue). Scale = 100 μm. **(B)** Percentage of GA_50_-GFP area that colocalized with antibody. N = 4 biological replicates from 2 independent experiments. Mean value of each replicate calculated from 3 distinct fields. **(C)** Flow cytometry-based approach to quantify the uptake of labeled antibodies into cells transfected with HA-GA_50_ or no plasmid (mock transfected). Membrane-associated antibodies were degraded by trypsin and any remaining extracellular signal was quenched by Trypan Blue. **(D)** HA-GA_50_ or mock transfected SH-SY5Y cells incubated with Alexa 488-labeled α-GA_b_ α-GA_3_ or the IgG control and analyzed by flow cytometry. Viable, singlet cells were selected for fluorescence negative (grey) or positive (orange, red or black) populations. **(E)** Percentage of antibody-positive cells after 48 hrs. Each data point represents 30,000 cells. N = 3 biological replicates. **(B, E)** Mean ± SD, one-way ANOVA followed by Tukey’s multiple comparison test. **(F)** Immunofluorescence of T98G cells expressing GA_161_-GA (green) and treated with antibody (red) for 72 hrs. **(G, H)** Compared with IgG control-treated cells, α-GA_1_-treated cells exhibited a significant reduction in the number of poly-GA aggregates per cell **(G)** and in volume of aggregates per cell **(H)**. N = 3 biological triplicates with 10 or 11 fields captured per replicate. Mean ± SD, unpaired two-tailed t-test. * P ≤ 0.05, ** P ≤ 0.01, *** P ≤ 0.001.

A flow cytometry-based approach using directly labeled human antibodies (**Fig. 1C**) was used to quantify antibody uptake by SH-SY5Y cells transfected to express HA-GA_50_ or exposed to transfection reagents without any plasmid (mock transfected). After incubation with fluorescently labeled α-GA_1_, α-GA_3_ or IgG isotype for 24 or 48 hrs, cells were treated with trypsin and trypan blue before analysis by flow cytometry to ensure that the detected fluorescence corresponds to internalized antibodies (36) (**Fig. 1 *C-E*** and *SI Appendix,* **Fig. S7*C-E***). As seen before with confocal microscopy (**Fig. 1*A*** and ***B*** and *SI Appendix,* **Fig. S7 *A*** and ***B***), accumulation of poly-GA increased the intracellular α-GA_1_ and α-GA_3_ antibody signal within 24 (*SI Appendix,* **Fig. S7*C***) or 48 hrs (**Fig. 1 *D*** and ***E***) compared to cells not expressing poly-GA (p < 0.001; **Fig 1*E*** and *SI Appendix,* **Fig. S7*C-E***). Only 2-6 % of the mock transfected (**Fig. 1*E***, *SI Appendix,* **Fig. S7*C***) or the non-transfected cells (*SI Appendix,* **Fig. S7 *D*** and ***E***) had detectable internalized antibodies while 15 to 22 % of poly-GA-transfected cells had internalized α-GA_1_ and α-GA_3_ antibodies. Notably, internalization of α-GA_3_ was more efficient than α-GA_1_ with a trend already observed after 24 hrs of treatment (p = 0.055; *SI Appendix,* **Fig. S7*C***) and a significant difference after 48 hrs (p < 0.001; **Fig. 1*E***). Detectable levels of internalized IgG isotype control were found in less than 9 % of the cells in all conditions.

To corroborate these findings, we used an independent, stable and inducible T98G glioblastoma cellular model overexpressing GA_161_-GFP. Following treatment with 100 nM α-GA_1_ or an IgG isotype control antibody for 72 hrs, cells were stained with anti-human IgG. Image analysis demonstrated that α-GA_1_ colocalized with poly-GA aggregates within T98G cells, while isotype control was not observed within cells (**Fig. 1*F***). Treatment of T98G cells with α-GA_1_ reduced the number of aggregates per cell by 29 % (**Fig. 1*G***) and reduced the volume of poly-GA aggregates per cell by 39 % (**Fig. 1*H***).

### Antibody uptake and colocalization with poly-GA in human neurons

The enhancement of antibody internalization or retention in presence of poly-GA was also seen in cultured human neurons. Neural stem cells (NSCs) were differentiated for 6 weeks into a functional neural network containing neurons and astrocytes (39), and treated for 72 hrs with ^ch^α-GA_3_ added to the medium. High-magnification confocal images revealed that ^ch^α-GA_3_ was internalized by human neurons (*SI Appendix,* **Fig. S8*A***, upper panels, arrowheads and inset) with no detectable signal in non-treated cells (*SI Appendix,* **Fig. S8*A***, lower panels). Neurons were transduced with a lentivirus expressing doxycycline-inducible GA_50_-GFP and treated for 24 hrs with ^ch^α-GA_1_, ^ch^α-GA_3_ or an IgG isotype control (**Fig. 2A** and *SI Appendix,* **Fig. S8*B***). Both ^ch^α-GA_3_ and the control antibodies were detected as extracellular clumps and internalized by GA_50_-GFP-expressing neurons at 72 hrs of treatment. While intracellular localization of ^ch^α-GA_3_ was observed in almost 100 % of GA_50_-GFP-expressing cells, the IgG isotype control antibody was rarely found accumulated intracellularly (18 %, p < 0.0001; **Fig. 2 *A*** and ***B***). In contrast with ^ch^α-GA_3_, intracellular ^ch^α-GA_1_ was detected only after 21 days of treatment (*SI Appendix*, **Fig. S8*B***). Three-dimensional reconstitution of confocal images of GA_50_-GFP-expressing human neurons treated with ^ch^α-GA_3_ revealed partial co-localization of GA_50_-GFP and ^ch^α-GA_3_ (*SI Appendix*, **Fig. S8*C***). Notably, we observed an incomplete colocalization of ^ch^α-GA antibodies with large round intracellular aggregates (*SI Appendix,* **Fig. S8*B***, inset), suggesting that the antibodies may not penetrate the dense core of these structures.

**Fig. 2.**
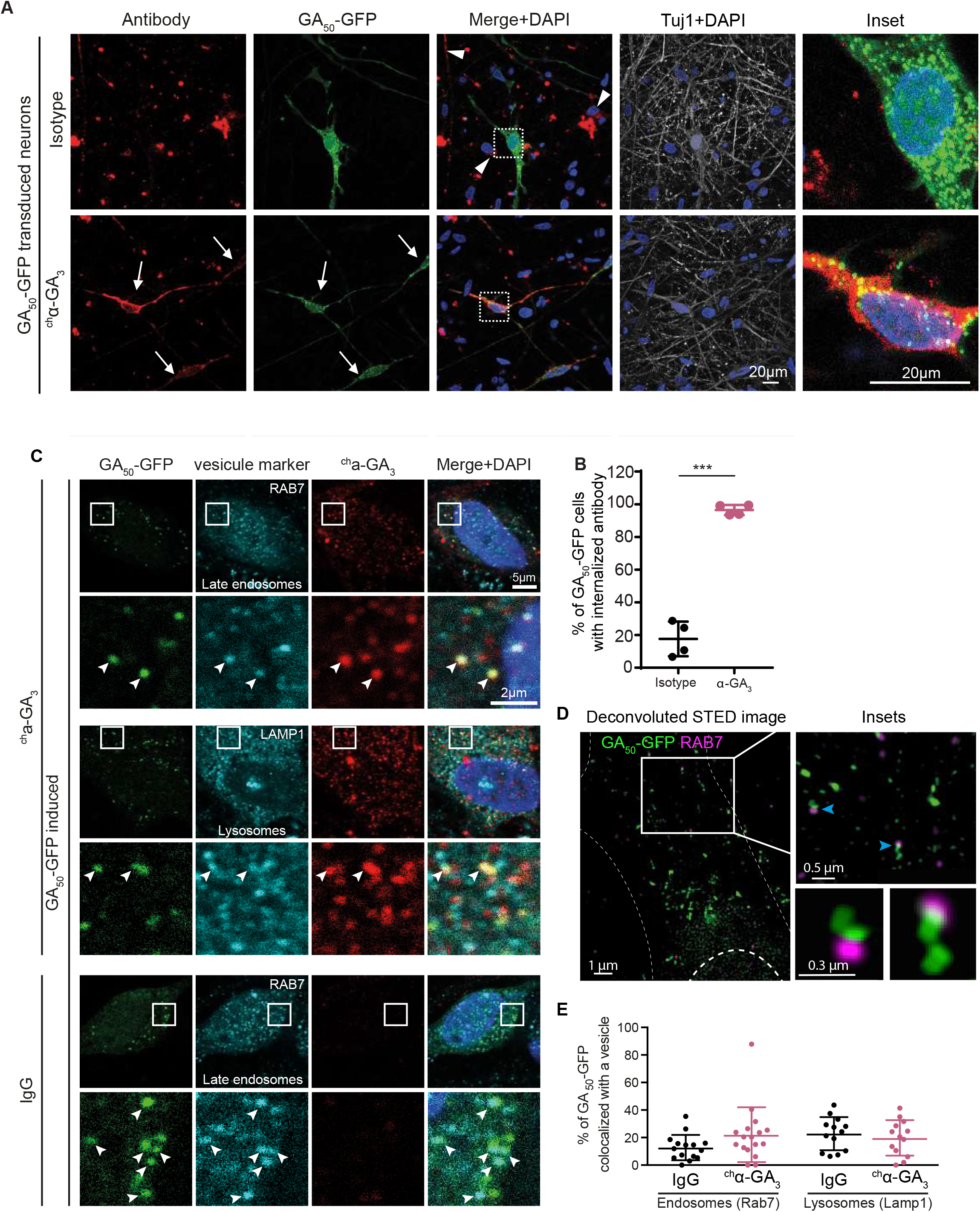
^ch^α-GA_3_ is internalized *via* vesicular compartments in human neurons, but does not alter GA_50_-GFP vesicular localization. **(A)** Confocal images of human neural cultures expressing an inducible GA_50_-GFP construct treated for 3 days with ^ch^α-GA_3_ or IgG control, and stained with a secondary α-human antibody (red). White arrows show antibody/GA_50_-GFP double positive cells and arrowheads show antibody uptake in cells not expressing GA_50_-GFP. Scale = 20 μm. **(B)** Quantification of the percentage of GA_50_-GFP expressing cells with internalized antibody. Four replicates, unpaired t-test. *** P < 0.0001. **(C)** Confocal images after staining with either α-RAB7 (late endosomes) or α-LAMP1 (lysosomes) and with α-mouse antibody (red) to detect the chimeric antibodies. **(D**) Deconvoluted STED image from a neuron expressing GA_50_-GFP treated with ^ch^α-GA_3_ and stained with RAB7. Inset of the STED image (upper panel) and the created 2D surface (lower panels) are enlarged from the original site (white box). (**E**) Distances between each GA_50_-GFP particle and the closest RAB7 or LAMP1 vesicular markers were measured. The percentage of GA_50_-GFP particles containing a vesicle within a 100 nm radius were considered as colocalized with the indicated vesicle. Each dot represents a field of view. Mann and Whitney U test (Rab7 p = 0.087 and Lamp1 p = 0.544).

To understand the intracellular compartment of the observed antibody-antigen interaction, we performed co-immunostaining of GA_50_-GFP, ^ch^α-GA_3_, and either RAB7 (endosomes) or LAMP1 (lysosomes). This analysis revealed that GA_50_-GFP and antibody partially colocalized with each of these markers (**Fig. 2*C***), supporting that intracellular interaction between poly-GA and externally delivered antibodies occurred within trafficking vesicles. Of note, the presence of the α-GA antibody was not required for the localization of GA_50_-GFP into endosomal vesicles (**Fig. 2*C***, lower panel). To determine if the presence of intracellular antibody faci litated the engulfment of GA_50_-GFP in intracellular vesicles, we quantified the colocalization between approximately 1400 GA_50_-GFP particles and each of these markers using super resolution microscopy (**Fig. 2 *D*** and ***E***). Antibody treatment with ^ch^α-GA_3_ showed a trend (p = 0.08) in favoring the colocalization of GA_50_-GFP with late endosomes compared to treatment with the IgG isotype control. Indeed 22 % of GA_50_-GFP vesicles were colocalized with Rab7 in cells treated with ^ch^α-GA_3_ compared to 12.7 % in cells treated with the IgG isotype control (**Fig. 2*E***). Localization of GA_50_-GFP in lysosomes was not affected (p = 0.54) (**Fig. 2*E***).

Taken together, these data demonstrate that α-GA antibodies entered cells and engaged intracellular poly-GA, the presence of which enhanced antibody uptake or intracellular retention. Antibody and GA_50_-GFP were found in similar intracellular vesicles, with no significant changes of GA_50_-GFP localization induced by ^ch^α-GA_3_ antibody treatment.

### Long-term antibody treatment in human neurons modulates poly-GA solubility by forming extracellular immune complexes

To determine whether chronic antibody treatment can modulate the aggregation state of poly-GA or trigger the aggregate clearance, we added ^ch^α-GA_1_, ^ch^α-GA_3_ or IgG isotype control antibodies to the culture medium of human neural culture transduced with inducible GA_50_-GFP. After 3, 7 or 21 days of poly-GA induction and simultaneous antibody treatment, cells were either fixed for immunofluorescence imaging followed by aggregate count or lysed for biochemical analysis (**Fig. 3*A***). Without antibody addition, only a faint, diffused or fine punctate GA_50_-GFP signal was detectable in the cytoplasm 3 days after induction (*SI Appendix,* **Fig. S9*A***). By 7 days, GA_50_-GFP either formed round and bright particles reminiscent of aggregates, or less bright and irregularly shaped structures resembling pre-inclusions (**Fig. 3*B***, *SI Appendix,* **Fig. S9*A***). At 21 days, both aggregates and pre-inclusions increased in number (**Fig 3*C***, *SI Appendix,* **Fig. S9*A***). Quantification confirmed the time-dependent increase of GA_50_-GFP intracellular inclusions (**Fig 3*C***, gray bars), which remained unaffected by the addition of ^ch^α-GA_1_ (**Fig. 3*C*)**, or ^ch^α-GA_3_ (*SI Appendix,* **Fig. S9*B***) into the cell culture medium.

**Fig. 3.**
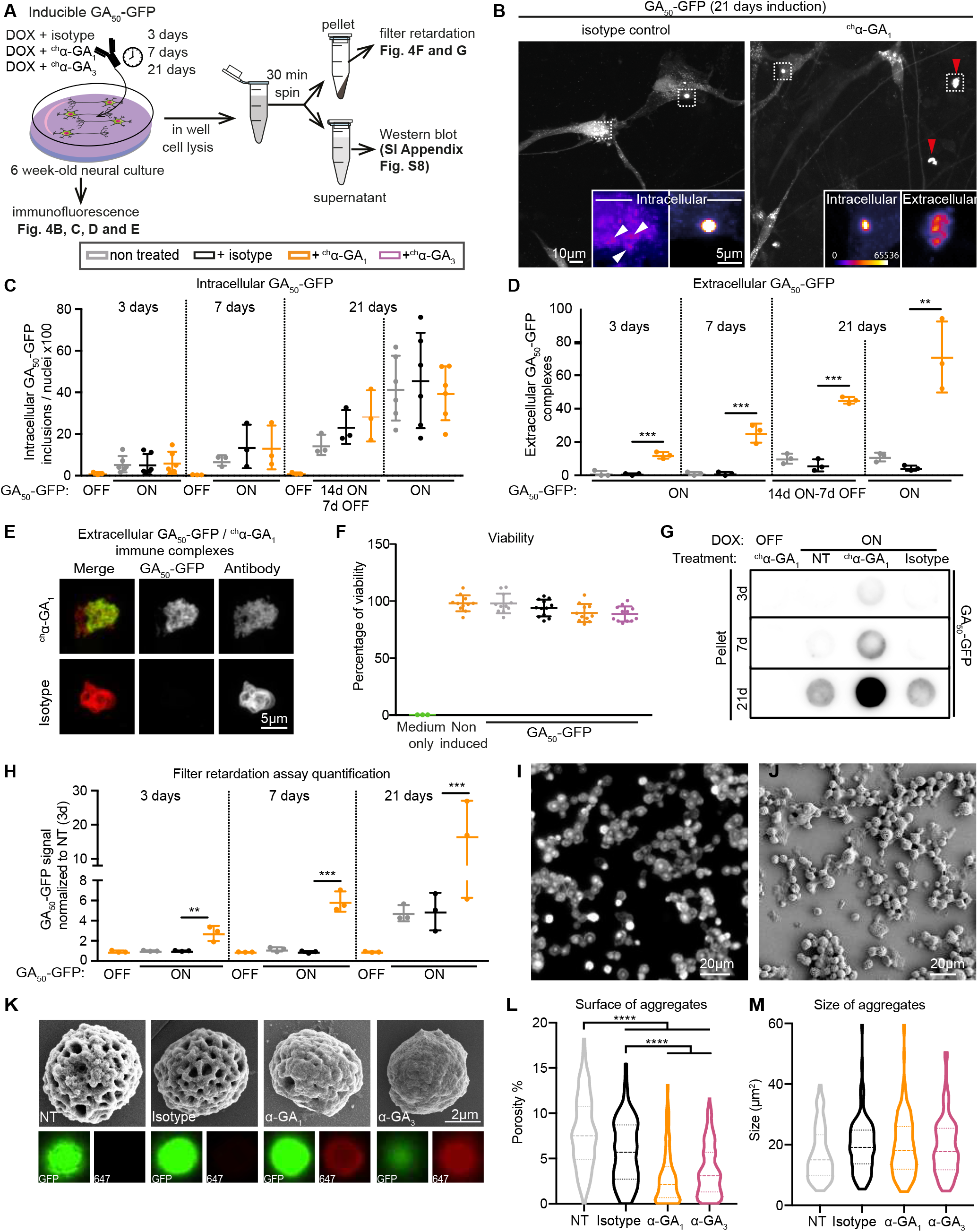
Poly-GA and antibodies form large hetero-complexes in long-term treated neuronal cultures. **(A)** Experimental strategies to test the effect of antibody treatment on human neurons expressing GA_50_-GFP. **(B)** Immunofluorescence showing GA_50_-GFP in neurons treated for 21 days with either an IgG control (upper panel) or ^ch^α-GA_1_ (lower panel). Large irregular extracellular GA_50_-GFP structures were observed only in samples treated with ^ch^α-GA_1_ antibody (red arrows and bottom right inset). Insets illustrate different GA_50_-GFP intracellular structures observed across all conditions including pre-inclusions (white arrows). Scale = 10 μm and 5 μm. **(C)** Quantification of intracellular GA_50_-GFP structures normalized to the number of nuclei, 36 images per well, 3-6 wells per condition. **(D)** Quantification of GA_50_-GFP extracellular structures normalized to the number of nuclei, 5 images per well, 3-6 wells per condition. Means +/− SD, beta-binomial test. **(E)** Confocal imaging of an ^ch^α-GA_1_ extracellular structure colocalizing with GA_50_-GFP (upper row), and an IgG control extracellular structure, not colocalizing with GA_50_-GFP (lower row). Antibodies were detected with α-mouse-Alexa 647. **(F)** Cell viability assay of human neurons expressing GA_50_-GFP treated with ^ch^α-GA_1_,_3_ or control antibody for 3 days. **(G)** Representative blots of a filter retardation assay where GA_50_-GFP was detected with an α-GFP antibody. **(H**) Quantification of the filter retardation blot for human neurons treated during 3, 7 or 21 days with ^ch^α-GA_1_ or IgG control. Intensity of each replicate was normalized to the 3 days IgG control samples. Unpaired t-test is used on the log10(x+1) transformed data. (**I**) Epifluorescent image of GA_50_-GFP aggregates isolated from transiently transfected HEK293T cells. (**J**) Isolated GA_50_-GFP aggregates visualized via SEM imaging. (**K**) Representative SEM and corresponding IF images of non-treated (NT) or Alexa Fluor 647-labelled isotype control, α-GA_1_- and α-GA_3_-treated GA_50_-GFP aggregates. (**L-M**) Quantification of the surface of the antibody treated GA_50_-GFP aggregates assessing their porosity (**L**) and the total area of the aggregates **(M)**. NT, n=139; isotype, n=125; α-GA_1_, n=183; α-GA_3_, n=193. Oneway ANOVA followed by Tukey’s multiple comparison test. P > 0.05 (no indication), ** P ≤ 0.01, *** P ≤ 0.001, **** P ≤ 0.0001.

Interestingly, the addition of ^ch^α-GA_1_ (**Fig. 3*B*** and ***D***) and ^ch^α-GA_3_ antibodies (*SI Appendix,* **Fig. S9*C***), but not of the IgG isotype control (**Fig. 3*B*** and ***D***), markedly increased the presence of extracellular bright, large, irregularly shaped GA_50_-GFP complexes (**Fig. 3*B***) which colocalized with ^ch^α-GA_1_ (**Fig. 4*E***) or ^ch^α-GA_3_ (SI Appendix, **Fig. S8*D***). Extracellular antibody-poly-GA complexes were stable for at least 7 days after doxycycline was removed to suppress new GA_50_-GFP production (**Fig. 3*D***). The formation of extracellular immune complexes was consistent with natural release of poly-GA from cells into the medium (34), as cell counting did not reveal poly-GA or antibody-mediated cell death (**Fig. 3*F***; *SI Appendix*, **Fig. S9*E***).

**Fig. 4.**
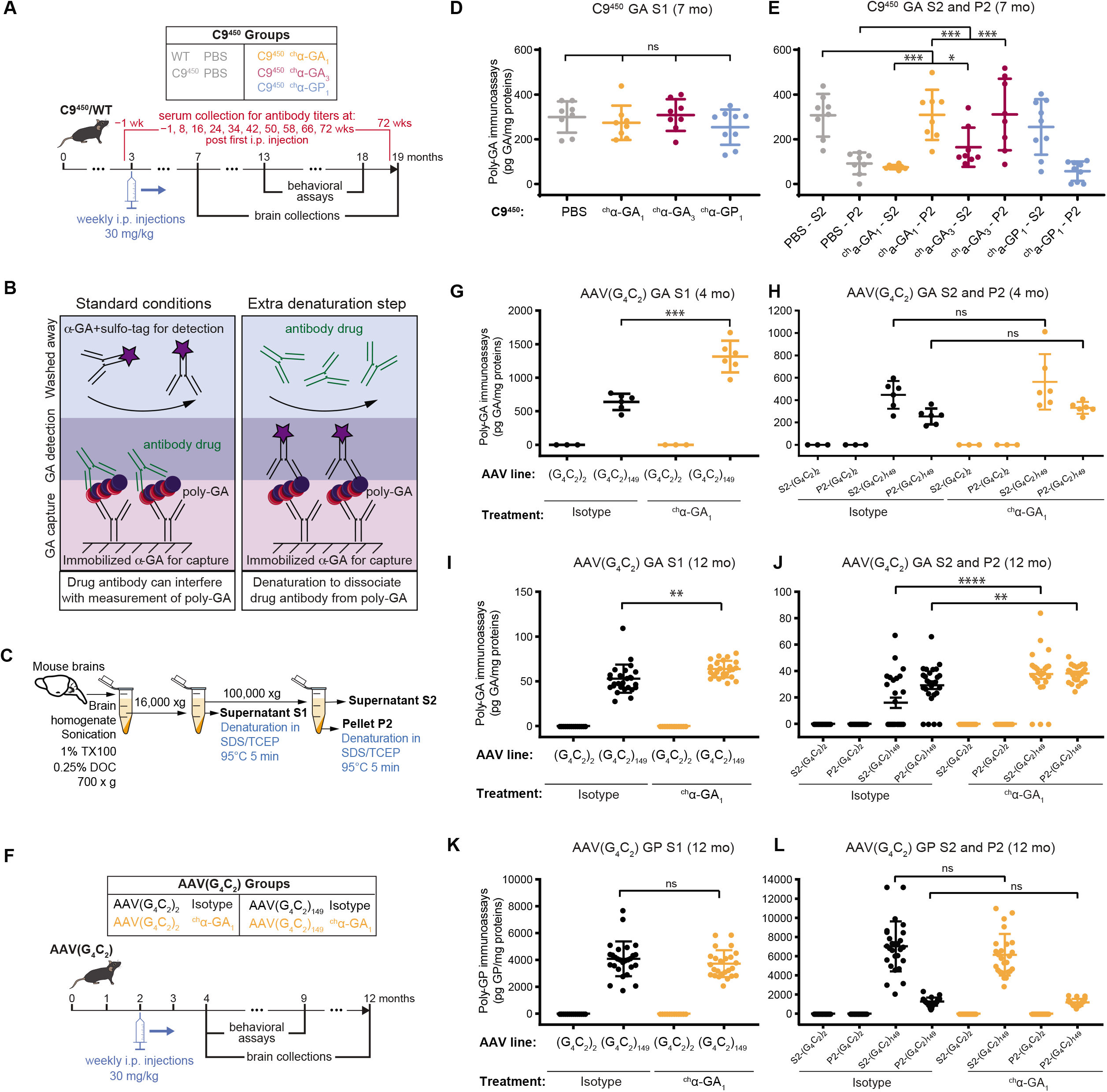
Immunoassay with sample denaturation identifies elevated levels of poly-GA in brains of mice treated with α-GA antibodies. **(A)** Scheme of chronic antibody treatment in C9^450^ mice receiving intra-peritoneal injection of PBS, ^ch^α-GA_1_, ^ch^α-GA_3_, or ^ch^α-GP_1_ antibodies from 3 to 19 months of age. **(B)** Scheme showing drug-antibody interference in the measurement of poly-GA protein levels using a sandwich-ELISA assay (left panel). Right panel illustrates the effect of sample denaturation prior to measurements. **(C)** Scheme of a mouse brain fractionation protocol adapted to denature samples to avoid interference of the antibody treatment by ELISA. Fractions highlighted in bold were analyzed. **(D, E)** Poly-GA levels measured by immunoassay from the supernatant S1 (**D**) and after ultracentrifugation (supernatant S2 and pellet P2) (**E**) of brains from 7-month-old C9^450^. **(F)** Scheme of chronic antibody treatment by intra-peritoneal injection of ^ch^α-GA_1_ or IgG control to AAV-(G_4_C_2_) mice from 2 to 12 months of age. **(G-J)** Poly-GA levels measured by immunoassay from the supernatant S1 (**G, I**) and after ultracentrifugation (supernatant S2 and pellet P2) (**H, J**) of brains from AAV-(G_4_C_2_) mice at 4 **(G, H)** and 12 **(I, J)** months of age. **(K, L)** Poly-GP levels measured by immunoassay from the supernatant S1 (**K**) and after ultracentrifugation (supernatant S2 and pellet P2) (**L**) of brains from AAV-(G_4_C_2_) mice at 12 months of age. Mean ± SD, one-way ANOVA followed by Tukey’s multiple comparison tests. Not significant (ns), P > 0.05, * P ≤ 0.05, ** P ≤ 0.01, *** P ≤ 0.001.

Biochemical analysis showed that the levels of soluble poly-GA were not different across experimental groups (*SI Appendix,* **Fig. S9*F***). On the other hand, poly-GA isolated by detergent solubilization followed by centrifugation (40) and quantified with a filter-retardation assay were markedly increased at all time points in neural cultures incubated with ^ch^α-GA_1_ (**Fig. 3*G*** and ***H***) or ^ch^α-GA_3_ (*SI Appendix,* **Fig. S9*G*** and ***H***) compared with control samples, supporting the presence of poly-GA/antibody immune complexes.

To test whether binding of α-GA antibodies to poly-GA aggregates alter their morphology, we first developed a biochemical method for the purification of GA_50_-GFP aggregates from transiently transfected HEK293T cells. Extracted GA_50_-GFP aggregates showed remarkable purity and homogeneity, permitting their characterization via scanning electron microscopy (SEM) (**Fig. 3*I***). GA_50_-GFP-expressing HEK293T cellular extracts were incubated with human Alexa Fluor 647-labelled α-GA_1_ α-GA_3_ or IgG isotype control antibodies, followed by poly-GA aggregate purification, brightfield, immunofluorescence and SEM imaging to eventually carry out correlative light-electron microscopy (CLEM) (**Fig. S10*A-N***). We then assessed and quantified the direct effects of antibody binding on the formation of poly-GA aggregates. Untreated or IgG isotype control-treated poly-GA aggregates appeared consistently spherical and with regular surface pores (**Fig. 3*K***). In contrast, binding of either α-GA_1_ or α-GA_3_ antibodies to poly-GA aggregates altered their morphology and yielded aggregates with a smoother surface (**Fig. 3*K-L***). Antibody binding had no effect on the size of the poly-GA spheres, which ranged roughly between 10 and 40 μm^2^ (**Fig. 3*M***). These results indicate that antibody binding directly alters the biochemical and potentially biological properties of GA_50_-GFP aggregates, which may affect their toxicity and spreading potential.

Overall, antibodies against poly-GA engaged extracellular GA_50_-GFP into detectable poly-GA/antibody immune complexes, without affecting soluble GA_50_-GFP levels or poly-GA intracellular structures, while they altered GA_50_-GFP aggregate formation and morphology in cellular extracts, potentially via stabilization of the poly-GA molecules within the aggregates.

### Pharmacokinetics and brain penetration of human-derived antibodies peripherally administered in C9^450^ mice

To test the therapeutic potential of antibodies with high affinity and specificity against poly-GA and poly-GP, three antibodies were selected for investigation in two different G_4_C_2_-expressing mouse models. The pharmacokinetic and antibody brain penetration properties of human-derived α-GA_1_, α-GA_3_ or α-GP1 antibodies were determined after a single intraperitoneal (i.p.) injection of 30 mg/kg of antibodies to transgenic mice expressing a human bacterial artificial chromosome (BAC) with 450 G_4_C_2_ repeats (C9^450^) (37) (*SI Appendix,* **Fig. S11*A-D***). The maximum concentrations (C_max_) in the plasma were 565±30, 338±24 and 587±54 μg/ml with estimated terminal elimination half-lives (t_1/2_) of 10.8, 11.0 and 7.2 days for α-GA_1_, α-GA_3_ and α-GP_1_, respectively. The corresponding C_max_ in the brain were 0.41±0.16, 0.12±0.07 and 0.28±0.09 μg/mg of total brain protein measured at 2 days post-injection. All antibodies were undetectable by three weeks post-administration. The ratio of the brain drug concentration to the plasma concentration measured at 2 days post-injection was of 0.05-0.1 %, consistent with previous reports for systemically administered antibodies (41). Immunofluorescence using human-specific IgG secondary antibodies did not detect α-GA_1_, α-GA_3_ and α-GP_1_ antibodies 10 days after a single i.p. injection in 20-month-old C9^450^ mouse brains having accumulated poly-GA aggregates (*SI Appendix,* **Fig. S11*E***).

### Chronic administration of human-derived antibodies modulates poly-GA solubility without significantly altering poly-GA levels in C9^450^ mice

To evaluate the effect of antibodies on the development of DPR pathology in mouse brain, antibodies were intraperitoneally injected in C9^450^ mice from 3 to 19 months of age (**Fig. 4*A***). To circumvent the mouse immune response towards the chronic administration of human antibodies we used murine IgG2a chimeric derivatives of the human antibodies. Chimeric α-GA antibodies (^ch^α-GA_1_, ^ch^α-GA_3_) and α-GP antibody (^ch^α-GP_1_) were administered once a week at 30 mg/kg in C9^450^ mice starting at 3 months of age (**Fig. 4*A***). The poly-GA aggregate load detectable by immunohistochemistry in C9^450^ mice was too low and variable in this cohort to be reliably quantified. Levels of soluble poly-GA and poly-GP were measured at 7 months of age by immunoassay after sonication of brain homogenates in the presence of 2 % SDS (37) (*SI Appendix,* **Fig. S12*A***, fraction 1) from C9^450^ mice expressing comparable levels of the transgene (*SI Appendix,* **Fig. S12*B***). Insoluble fractions obtained by ultracentrifugation and resuspension of the corresponding pellet in 7 M Urea (*SI Appendix,* **Fig. S12*A***, fraction 2) were measured using similar immunoassays. Soluble poly-GP did not significantly differ between treatment groups (*SI Appendix,* **Fig. S12*C***). Poly-GA proteins, however, were not detectable in mice treated with ^ch^α-GA_1_ and ^ch^α-GA_3_ and were significantly reduced in mice treated with ^ch^α-GP1 (an antibody recognizing poly-GA with lower affinity; *SI Appendix,* **Fig. S1*A*** and **S4*B***) compared to mice injected with saline only (*SI Appendix,* **Fig. S12*D***). This observation suggests that α-GA antibodies interfere with the detection of poly-GA in immunoassays that do not include denaturation of the samples, likely by masking of the epitopes by the injected antibody (**Fig. 4*B***, left panel). In addition, while poly-GA was normally not detected in the urea-insoluble fraction from saline injected C9^450^ mice (*SI Appendix,* **Fig. S12*E*,** C9^450^ PBS), poly-GA was present in the insoluble fraction from mice treated with ^ch^α-GA_1_ or ^ch^α-GA_3_ antibodies (*SI Appendix,* **Fig. S12*E***, fraction 2).

Notably, similar results were obtained when antibodies were directly spiked into mouse brain homogenates further confirming that poly-GA antibodies form immune complexes with poly-GA that migrate in insoluble fractions and interfere with immunoassay’s detection when samples are not efficiently denatured (*SI Appendix,* **Fig. S12*F-H***).

To accurately investigate the effect of antibody treatment on poly-GA levels, we adapted the protocol by denaturing any carry-over antibody that might interfere with the poly-GA immunoassay using resuspension in SDS/tris(2-carboxyethyl)phosphine (TCEP) and boiling of the samples (**Fig. 4*B***, right panel, and **Fig. 4*C***). The samples were also subjected to centrifugation and ultracentrifugation after homogenization in 1 % TX100 and 0.25 % deoxycholate (DOC) (**Fig. 4*C***) (1). Denaturation of the brain homogenates demonstrated that poly-GA levels were unchanged between the different treatment groups in the first supernatant fraction (S1) (**Fig. 4*D***). Consistent with our previous results (*SI Appendix,* **Fig. S12*E***, fraction 2), both ^ch^α-GA_1_ and ^ch^α-GA_3_ i.p. injections in C9^450^ mice increased the presence of poly-GA in the pellet fraction (P2) after ultracentrifugation (**Fig. 4E**), supporting the presence of poly-GA/antibody immune complexes.

### Chronic administration of human-derived antibodies modulates poly-GA solubility and increases poly-GA levels in AAV(G_4_C_2_)_449_ mice

The impact of ^ch^α-GA_1_ antibody was also determined in somatic transgenic mice generated by intra-cerebroventricular (ICV) administration to post-natal day 0 mice of adeno-associated virus encoding either 2 or 149 G_4_C_2_ hexanucleotide repeats [AAV(G_4_C_2_)_2_ or AAV(G_4_C_2_)_149_] (**Fig. 4*F***). Weekly i.p. injections of ^ch^α-GA_1_ or the IgG isotype control were carried out from 2 to 12 months of age and brains were collected either at 4 or at 12 months of age for poly-GA and poly-GP measurements (**Fig. 4*G-L***). Using an immunoassay that included denaturation of the samples, we identified a significantly increased accumulation of poly-GA in the supernatant S1 fraction in AAV(G_4_C_2_)_149_ mice treated with ^ch^α-GA_1_ compared to mice injected with the IgG control at 4 (**Fig. 4*G***) and 12 months (**Fig. 4*I***). After ultracentrifugation of the samples, the levels of poly-GA in protein fractions S2 and P2 were not changed at 4 months of age (**Fig. 4*H***), but were significantly increased in all fractions from 12-month-old mice treated with ^ch^α-GA_1_ compared to mice injected with the IgG control (**Fig. 4*J***). On the contrary, poly-GP solubility and levels were unaltered by ^ch^α-GA_1_ treatment (**Fig. 4*K*** and ***L***).

In addition, sarkosyl-insoluble pellets isolated via SarkoSpin (40) from total brain homogenates of 4-month-old AAV(G_4_C_2_) mice were analyzed via filter retardation assay (*SI Appendix,* **Fig. S13*A*** and ***B***). ^ch^α-GA_1_ antibody was specifically retained on the membrane in the ^ch^α-GA_1_-treated AAV(G_4_C_2_)_149_ mouse samples suggesting that the non-denaturing conditions of the SarkoSpin protocol led to the isolation of sarkosyl-insoluble ^ch^α-GA_1_ antibody-poly-GA complexes, which were not observed with IgG isotype control (*SI Appendix,* **Fig. S13*A*** and ***B***). Insoluble, poly-ubiquitinated proteins were detected in both AAV(G_4_C_2_)_149_ mouse conditions and their levels were not affected by ^ch^α-GA_1_ antibody treatment (*SI Appendix,* **Fig. S13*C*** and ***D***).

### Chronic administration of human-derived antibodies did not impact poly-GA aggregate load in AAV(G_4_C_2_)ι_49_ mice

By 4 months of age, AAV(G_4_C_2_)_149_ mice accumulated large perinuclear poly-GA aggregates throughout the brain that co-localized with poly-GR and poly-GP (*SI Appendix,* **Fig. S14*A*** and ***B***), as observed in postmortem tissues from patients (7). We determined the area occupied by poly-GA aggregates in AAV(G_4_C_2_)_149_ mice treated for 2 or 10 months with ^ch^α-GA_1_ antibody compared to mice treated with the IgG isotype control. To test whether treatment with α-GA antibodies may interfere with detection of aggregates (as observed in immunoassays without strong denaturation; **Fig. 4*B***, *SI Appendix,* **Fig. S12*D*** and ***G***), immunofluorescence was performed using either an antibody raised against poly-GA (37) or an antibody raised against a N-terminal peptide starting at a CUG initiation codon in the poly-GA frame (10). When using an anti-GA antibody to detect aggregates there was no change in the poly-GA aggregates after 2 months of treatment (*SI Appendix,* **Fig. S14*C*** and ***D***), but the area appeared significantly decreased in the cortex after 10 months of treatment with ^ch^α-GA_1_ antibody (*SI Appendix,* **Fig. S14*E***). A non-significant similar trend was observed in the hippocampus (*SI Appendix,* **Fig. S14*F***). However, this reduction was not observed when we used an antibody raised against the N-terminal peptide of poly-GA (**Fig. 5*A-C***), demonstrating the importance of using antibodies that recognize different epitopes than the treatment antibody when assessing the effect of an immunotherapy against poly-GA.

**Fig. 5.**
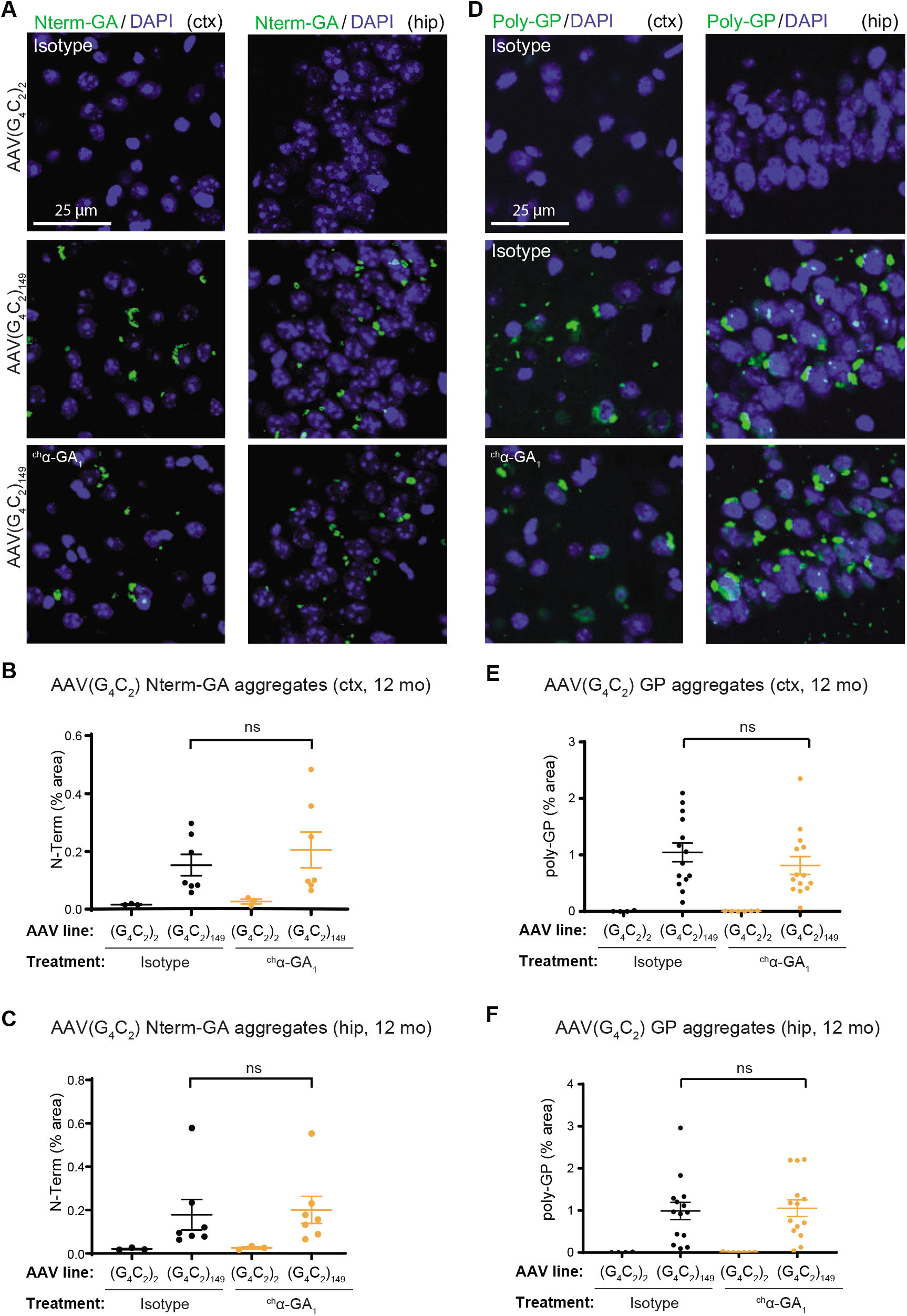
Chronic administration of human-derived α-GA_1_ antibody does not reduce the poly-GA and poly-GP aggregate load in brains of AAV(G_4_C_2_)_149_ mice. **(A)** Immunofluorescence of poly-GA staining in the motor cortex (left) and hippocampus (right) of 12-month-old AAV-(G_4_C_2_)_2_ or AAV-(G_4_C_2_)_149_ mice treated with ^ch^α-GA_1_ or an IgG control. **(B-C)** Quantifications of percent area occupied by poly-GA aggregates detected with a N-terminal-poly-GA antibody in cortex **(B)** and hippocampus **(C)**. (**D**) Immunofluorescence of poly-GP staining in the motor cortex (left) and hippocampus (right) of 12-month-old AAV-(G_4_C_2_) mice treated with ^ch^α-GA_1_ or an IgG control. **(E-F)** Quantifications of percent area occupied by poly-GP staining in cortex **(E)** and hippocampus **(F)**. Scale = 25 μm. Mean ± SD, one-way ANOVA followed by Tukey’s multiple comparison test. Not significant (ns), P > 0.05, * P ≤ 0.05.

As expected, poly-GP aggregates were not affected in either the cortex (**Fig. 5*D*** left panels, and **Fig. 5*E*)** or the hippocampus (**Fig. 5*D*** right panels, and **Fig. 5*F***). The level of poly-GR measured by immunoassay (*SI Appendix,* **Fig. S14*G***) and the area of poly-GR aggregates (*SI Appendix,* **Fig. S14*H***) were also not modified by treatment with ^ch^α-GA_1_ antibody. Similarly, the number of phospho-TDP-43 aggregates detected by immunohistochemistry in AAV(G_4_C_2_)_149_ mice was not altered by treatment with ^ch^α-GA_1_ (*SI Appendix,* **Fig. S15*A*** and ***B***).

### Long-term *in vivo* administration of DPR antibodies was well tolerated with a modest impact on behavior in C9^450^ mice

C9^450^ mice were treated by weekly injection of ^ch^α-GA_1_, ^ch^α-GA_3_ and ^ch^α-GP_1_ from 3 to 19 months of age (**Fig. 4A**). Antibody titers in serum (measured every 2 months, 24 hrs after injection) remained stable over time (*SI Appendix,* **Fig. S16*A-C***). Chronic administration did not result in any obvious adverse effects, with comparable survival (**Fig. 6A**) and body weight (*SI Appendix,* **Fig. S16*D*** and ***E***) between the different treatment groups, demonstrating the tolerability of all three antibodies at 30 mg/kg per week for 16 months. At 13 months of age, only C9^450^ males exhibited a decreased activity with a significant reduction in distance moved compared to age-matched wild-type mice in an open-field assay (*SI Appendix,* **Fig. S16*F***). At this age, these differences were not impacted by ^ch^α-GA_1_, ^ch^α-GA_3_ and ^ch^α-GP_1_ antibody treatment (*SI Appendix,* **Fig. S16*F*)**. However, by 18 months of age, C9^450^ mice treated with PBS continued to display a significantly decreased activity compared to wild-type animals, while mice treated with ^ch^α-GA_3_ showed a significant rescue when compared to C9^450^ animals treated with PBS (p=0.0149) (**Fig. 6*B***). Mice treated with ^ch^α-GA_1_ and ^ch^α-GP1 antibodies also showed a non-significant trend towards improvement in this behavioral assay (**Fig. 6*B***). As previously reported (37), C9^450^ mice develop a loss of hippocampal neurons that was not significantly alleviated by treatment with DPR antibodies (**Fig. 6*C***).

**Fig. 6.**
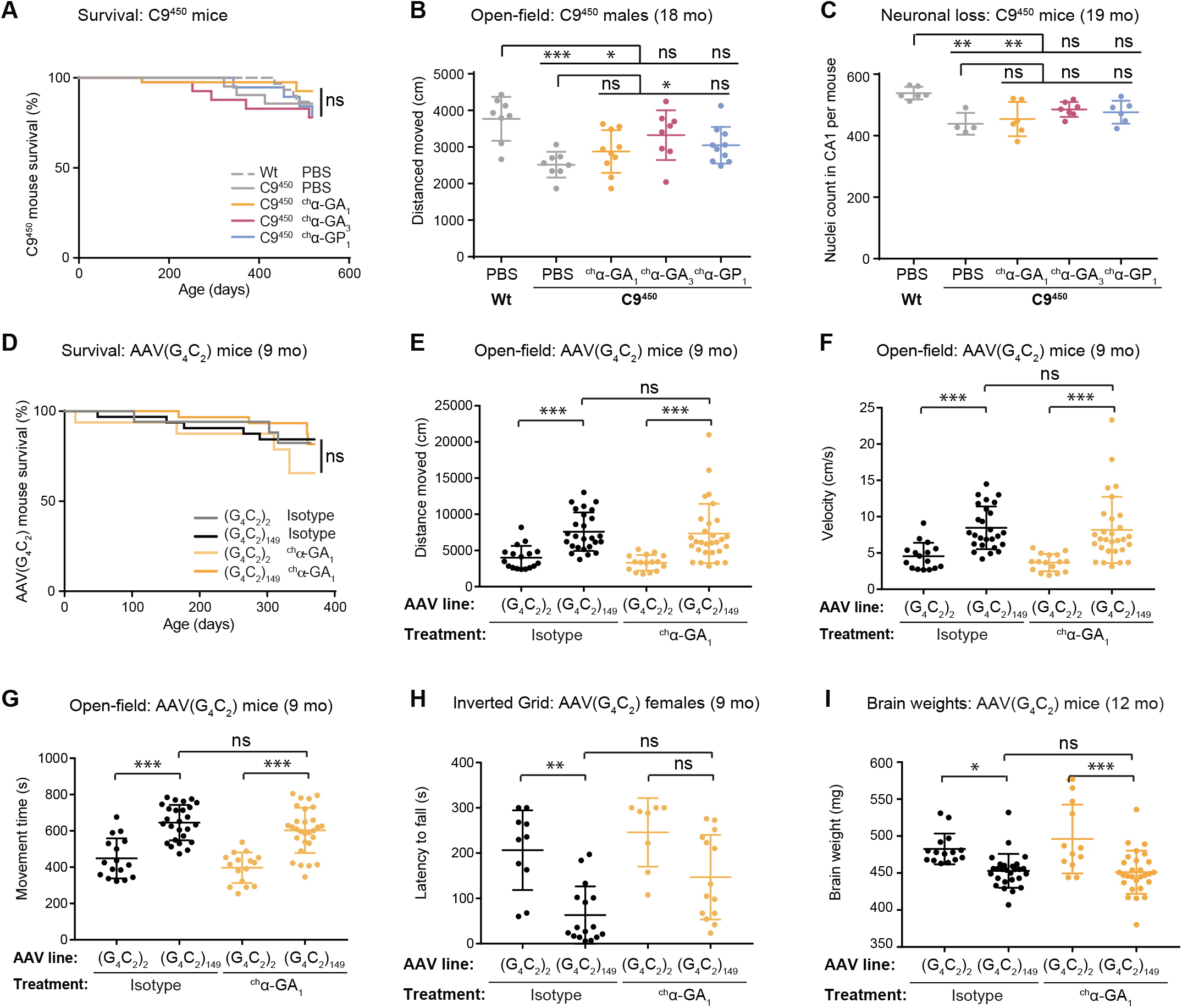
Chronic administration of α-GA antibodies impacted only a subset of behavioral and neurodegeneration phenotypes in one of two C9ORF72 mouse models. **(A)** Survival Kaplan–Meier curves of C9^450^ and wild-type mice receiving injections of PBS, ^ch^α-GA_1_, ^ch^α-GA_3_ and ^ch^α-GP_1_ antibodies for 16 months. **(B)** Distance traveled in the openfield test by 18-month-old males (n ≥ 8 per group). Mean ± SD, Kruskal-Wallis test followed by Dunnett’s multiple comparison tests. **(C)** Nuclei quantification in the hippocampal CA1 region in 19-month-old mice (n ≥ 4 mice per group; n ≥ 3 matched sections per mouse). Mean ± SD, one-way ANOVA followed by Dunnett’s multiple comparison tests. **(D)** Survival Kaplan–Meier curve of AAV-(G_4_C_2_)_2_ and AAV-(G_4_C_2_)_149_ mice receiving injections of ^ch^α-GA_1_ or IgG isotype control for 10 months. **(E-G)** Distance traveled **(E)**, velocity of movement **(F)** and time spent moving **(G)** in the open-field test. **(H)** Time taken to fall from inverted grid by 9-month-old AAV-(G_4_C_2_) female mice. **(I)** Brain weights of AAV-(G_4_C_2_) mice treated for 10 months. Mean ± SD, one-way ANOVA followed by Dunnett’s multiple comparison tests. Not significant (ns), P > 0.05, * P ≤ 0.05, ** P ≤ 0.01 and *** P ≤ 0.001.

### Long-term *in vivo* administration of DPR antibodies did not significantly impact disease progression in AAV(G_4_C_2_)_149_ mice

AAV(G_4_C_2_)_149_ mice were treated by weekly i.p. injection of ^ch^α-GA_1_ at 30 mg/kg from 2 to 12 months of age (**Fig. 4*E***). The survival of these mice was not impacted by the expression of the G_4_C_2_ repeats as previously described (42) and antibody treatment was well tolerated (**Fig. 6*D*** and *SI Appendix,* **Fig. S16*G-H***). AAV(G_4_C_2_)_149_ mice presented abnormal activity including increases in distance traveled and velocity of movement, and time spent moving on an open-field assay compared to control AAV(G_4_C_2_)_2_ mice (**Fig. 6*E-G***). These phenotypes were not impacted by chronic administration of ^ch^α-GA_1_ antibody. When assessing strength by measuring the ability to cling on an inverted metal grid, female AAV(G_4_C_2_)_149_ mice showed significantly lower performance compared to AAV(G_4_C_2_)_2_ mice (p = 0.002) (**Fig. 6*H***). Although there was a trend towards improvement, the deficit in AAV(G_4_C_2_)_149_ mice was not significantly rescued by ^ch^α-GA_1_ treatment (p = 0.11). In addition, treatment with ^ch^α-GA_1_ did not impact the decrease in brain weight observed in AAV(G_4_C_2_)_149_ mice compared to AAV(G_4_C_2_)_2_ mice (**Fig. 6I**).

## Discussion

In this study, we have characterized potential immunotherapies (based on human antibodies) for C9ORF72-related ALS and FTD. Eleven antibodies against all five C9ORF72 DPR species were identified and systematically characterized. The exact trigger(s) leading to the production of antibodies against DPRs in healthy people is unknown. It is conceivable that due to their highly repetitive sequences, DPRs may present sequence or structural similarities with other antigens potentially derived from bacteria or viruses. Alternatively, the 2 to 30 (G_4_C_2_) repeats found in the normal population may produce DPR proteins at a very low rate, which may trigger an antibody-mediated immune response without being pathogenic. While we have systematically characterized antibodies against each C9ORF72-related DPR, we focused on antibodies against poly-GA, recognizing that this DPR is the most abundant with high aggregation propensity in ALS/FTD human autopsy brain samples (7, 43) and with strong neurotoxicity in mice (20, 23). Moreover, poly-GA has the ability to trap other DPR species and modulate C9ORF72 toxicity observed in multiple cellular and animal models (21, 22, 24).

While antibody treatment against intracellular Tau and α-synuclein are currently being tested for Alzheimer’s and Parkinson’s diseases respectively (28, 29, 33), the exact mechanisms of action of immunotherapy against intracellular proteins remain unclear. Antibodies were shown to either facilitate clearance of the target or to prevent spreading and toxicity. The tested α-GA antibodies were robustly internalized by poly-GA-expressing human neurons, a finding in line with published studies showing antibody uptake by neuronal cells (35, 36). In this cellular model, intracellular antibodies colocalized with poly-GA in cytoplasmic puncta, however, less prominently to very dense and large intracellular aggregates formed over time. It is possible that their compact structure may conceal the epitope recognized by the antibody. Alternatively, poly-GA physical associations with other proteins may interfere with epitope recognition. Antibody and poly-GA colocalized partially with late-endosomes and lysosomes, but anti-GA antibody treatment did not significantly alter GA_50_-GFP vesicular localization. Whether antibody and antigen entered the same degradation pathway independently, or antibody binding on poly-GA triggered its engulfment in endocytic vesicles, thereby potentially stimulating its clearance remains unanswered. It is also possible that the antibodies engaged the poly-GA extracellularly and entered human neurons already as a complex. However, the absence of antibody-engaged poly-GA in non-transduced, wild-type neurons present in the same neuronal network challenges that notion. Rather, as supported by super resolution microscopy, the dense GA_50_-GFP inclusions and aggregates may be present in a different compartment than the antibody-engaged poly-GA. Notably, despite target engagement in our cellular models and reduction of poly-GA aggregates in T98G anchorage independent cancer cell line, intracellular GA_50_-GFP inclusions and aggregates were not affected by antibody treatment in cultured human neurons, highlighting differences between cell types.

Of note, the three cellular models used in this study were differently modified to overexpress poly-GA. While the liposome-mediated transfection of SH-SY5Y cells may have indirectly enhanced the antibody “uptake” because of partially compromised cell membrane, both the T98G cells (stable transfection) and human neurons (lentivirus-mediated gene delivery and cell recovery for several days before antibody treatment) likely had intact cell membrane and thus accurately modeled antibody uptake and/or retention in cancer cells or human neurons, respectively. Interestingly, anti-GA antibodies did engage less prominently the dense poly-GA aggregates in human neurons synthetizing poly-GA, compared to their targeting of aggregates in stably transfected T98G cancer cells. This points to a distinct aggregate handling between cycling cells and differentiated neurons. In addition, IF experiments with fixed and permeabilized motor neuron-like NSC-34 cells overexpressing poly-GA via transient transfection revealed that all tested anti-GA antibodies (α-GA_1-4_) only partially recognized dense poly-GA aggregates.

It was previously shown that poly-GA is released into the extracellular space and can be taken up by neighboring cells, thereby increasing DPR aggregation (22, 32, 34). A similar mechanism may account for spreading of DPR pathology throughout the nervous system, as was hypothesized for other intracellular protein aggregates found in ALS/FTD (44), a process that may be blocked by immunotherapy. We showed that human-derived antibodies efficiently captured extracellular poly-GA over the course of 3 weeks forming large immune complexes in human neuronal cultures. This was also described in the context of Alzheimer’s disease, where an anti-Tau antibody blocked toxicity and spreading through the formation of immune complexes (45). We were unable to assess whether the formation of poly-GA/antibody immune complexes could rescue poly-GA toxicity in this cellular system, since we found no detectable poly-GA toxicity within the time course of 21 days.

In contrast to previous studies that reported a decrease in intracellular aggregates and insolubility of poly-GA upon α-GA treatment in cells (1, 32), our analysis in cultured human neurons did not find a decrease of intracellular inclusions and showed a significant increase of poly-GA insolubility, likely due to the formation of immune complexes (**Fig. 3**). Antibody-induced insolubility has been previously reported for α-synuclein, which formed amorphous aggregates *in vitro* in the presence of four out of six tested α-synuclein antibodies (46). In comparison to α-synuclein, which has a strong ability to form fibrils (46), the unusually high hydrophobic and low complexity nature of poly-GA (7, 15) may make it more prone to clump into amorphous aggregates when molecules are brought into close proximity following antibody binding. An antibody selective for soluble Tau triggered the formation of extracellular complexes and protected against exogenous paired helical filament toxicity, while an alternative antibody directed against aggregated Tau failed at forming extracellular immune complexes and could not confer cellular protection (45).

We also observed that the detection of poly-GA either by immunoassay or by immunofluorescence staining was altered by treatment with anti-GA antibodies. Indeed, using an antibody that does not recognize the poly-GA epitopes but rather a N-terminal peptide translated in frame with poly-GA (10), we demonstrated that poly-GA aggregates were not affected by 9 months of ^ch^α-GA_1_ treatment in AAV(G_4_C_2_)_149_ mice (**Fig. 5**). However, the aggregate load appeared reduced when immunostaining was performed using an antibody against poly-GA (*SI Appendix,* **Fig. S14*E***), suggesting that the treatment with ^ch^α-GA_1_ antibody may block the recognition of poly-GA epitopes by the detecting antibody leading to underestimation of poly-GA aggregates. The vast majority, if not all, of poly-GA is translated from a start codon located 24 nucleotides upstream of the repeat that encode the N-terminal peptide (10, 47). Hence, it is unlikely that the poly-GA species detected by the N-terminal peptide directed-antibody represent only a subset of poly-GA that would be differently impacted by the treatment. In addition, an interference between the treatment and detection antibodies was demonstrated in a biochemical assay lacking efficient denaturation of the samples before poly-GA measurement (*SI Appendix,* **Fig. S12**). Such an interference was not observed for the uncharged, flexible and highly soluble (48) poly-GP molecules (*SI Appendix,* **Fig. S12*C***) suggesting that intrinsic structural features of poly-GA are altered by antibody recognition, as supported by our SEM analysis of purified poly-GA (**Fig. 3*I-M***). Combined with the robust formation of poly-GA-antibody complexes evident in all our cellular work, these changes highlight the necessity of analyzing all biochemical fractions when comparing antibody-treated to non-treated conditions. While interference between the treatment and detecting antibodies or antibody-induced biochemical changes may not be an issue for all proteins, our study demonstrates that careful denaturation of biochemical samples and use of antibodies recognizing independent epitopes is warranted for accurate assessment of the impact of immunotherapies on aggregation-prone proteins.

In this study, we have not observed a reversal of the clinical phenotypes linked to C9ORF72 disease in our antibody-treated AAV(G_4_C_2_)_149_ mice. This is contrary to a recently published study in a C9ORF72 BAC model (1). The reason for this discrepancy in response to the antibody treatment is unknown ‖ one possibility is that the different mouse models used in the two studies display different phenotypes resulting in differential responses to antibody treatment. Alternatively, the discrepancy may be linked to the fact that in contrast to other described mouse models expressing G_4_C_2_ C9ORF72 repeats (37, 38, 42, 49, 50), the model used in the Nguyen et al. study has been reported to develop severe neurodegeneration (1). Mordes et al (51) described that two independent cohorts of the same model had similar levels of DPRs but did not have the same behavioral phenotypes previously reported in these mice. While independent laboratories reproduced the originally reported phenotypes in the C9ORF72 BAC mouse model (52), Mordes and colleagues (51) proposed that the severe neurodegeneration reported in a fraction of the mice used by Nguyen and colleagues may be linked to the space cadet syndrome (SCS), previously reported in WT mice with an FVB/N background (53). While C9ORF72-linked neurodegeneration may be exacerbated by – and potentially distinguished from – the severe seizure phenotypes affecting both C9ORF72 and non-transgenic animals (52), future studies are necessary to determine the relative contribution(s) of the *C9ORF72* repeat expansion, the DPR expression levels and the SCS-linked pathologies to the described phenotypes and their immunotherapy-driven reversal in different mouse models.

In the current study, we have not observed target engagement by IHC in the brains of our C9^450^ antibody-treated mice. We have observed constant antibody plasma titers over 1.5 years with brain penetration of a small fraction (~0.1 %) of the injected antibody (as previously shown (25) for other antibodies), but peripherally injected human antibodies were not found to co-localize with neuronal poly-GA aggregates (*SI Appendix,* **Fig. S11*E***). The reason for this difference from the observations in Nguyen et al. (1) is unclear, but it is conceivable that the severe neurodegeneration and/or SCS pathology in the mice used by Nguyen et al. may be associated with blood-brain barrier leakage, which might facilitate antibody entry to the brain. Whether that accounts for the reported decrease in poly-GA load in these mice remains to be clarified.

In our C9^450^ cohort, the levels of poly-GA and poly-GP could be measured by immunoassay but the number of poly-GA aggregates were too low to evaluate the effect of antibody treatment on poly-GA aggregate load. Despite the lack of widespread poly-GA pathology, long term treatment with anti-GA antibodies improved an open field movement test in aged C9^450^ mice (albeit modestly). In AAV(G_4_C_2_) 149 mice, anti-GA treatment failed to ameliorate brain atrophy and poly-GA levels increased following treatment during 9 months. This finding supports that at least a small portion of peripherally injected antibodies accessed poly-GA in the brain and impacted its turn-over. Targeting some of the other reportedly toxic DPR proteins, such as poly-GR and poly-PR, may be an alternative and potentially synergistic approach to treat *C9orf72* disease which should be explored in future studies. Furthermore, we anticipate that antibody delivery is key for the success of immunotherapy and approaches to increase antibody penetration to the central nervous system (54) might result in enhanced therapeutic benefit.

## Acknowledgments

We are grateful to Moritz Kirshmann and Joe Weber for support in data analysis, to Ulrich Wagner for statistical analysis, to Nicola Bothwick for mouse brain analysis, and to Moaz Abdelrehim and Mohamed Al Abdulla for technical help. We are grateful to Asvin Lankkaraju in Adriano Aguzzi’s laboratory for providing Rab7 and Lamp1 antibodies. We thank all the members of the Albers, Wainger and Aguzzi laboratories for helpful discussions. Imaging was performed at the Center for Microscopy and Image Analysis with assistance from Johannes Riemann and Andres Kaech (University of Zurich) as well as with microscopes from Rudy Tanzi’s and Brian Wainger’s laboratories (MGH). Human postmortem tissues were obtained from Matthew Frosch and Jose McLean at the Massachusetts Alzheimer’s Disease Research Center (PG50 AG005134).

## Author contribution

RT, CZL, KDM, MJ, MHP, ARS, NM, CA, PB, KS, NM, NC, PB, SD, ID, CCL, JJ, XJ, JP, SN, YG, SD, IDL, ZM, JW, MW, MM, KS, TG, MP, FM and CLT conducted experiments or analyzed data; KJW, JJ, FF, NL, PDR, MPB, LMD, EB, YZ, JB, BW, MWK, MR, CH, RMN, DWC, LP supplied material and/or gave significant input; MJ, KDM, RT, MHP, ARS, MWK, MR, MP, CLT, FM, JG, DWC, LP wrote or edited the manuscript.

## Data and materials availabilities

The data that support the findings of this study are available from the corresponding authors upon request. Constructs and reagents generated in this study can be shared upon request with establishment of an appropriate material transfer agreement. There may be restrictions to the availability of some data/reagents due to agreements with industry partners.

