## Supplementary Material for "Comprehensive preclinical evaluation of human-derived anti-poly-GA antibodies in cellular and animal models of C9ORF72 disease"

**This PDF file includes:**

Materials and Methods
Table S1, Figs. S1 to S16

### Materials and Methods

*Production of recombinant human antibodies*

Human DPR-protein-targeting antibodies were derived from a blood lymphocyte library collected from healthy elderly subjects as previously described (25). In brief, memory B cells were isolated by anti-CD22-mediated sorting from peripheral blood lymphocyte preparations. Isolated cells were cultured on gamma-irradiated human peripheral blood mononuclear cell feeder layers. Supernatants from isolated human memory B cells were screened for binding to $GA_{15}$, $GP_{15}$, $GR_{15}$, $PA_{15}$ or $PR_{15}$ synthetic DPR peptides (see below). Positive antibody hits were cloned using cDNA of IgG heavy and kappa or lambda light chain variable region sequences. Selected antibodies were then sub-cloned in expression constructs using Ig-framework specific primers for human variable heavy and light chain families in combination with human J- and H- segment-specific primers. All presented antibodies were engineered to incorporate glycosylated human IgG1 heavy and human lambda light chain constant domain sequences. Murine chimeric IgG2a/lambda version of α-$GA_1$, α-$GA_3$, and α-$GP_1$ ($^{ch}$α-$GA_1$, $^{ch}$α-$GA_3$, and $^{ch}$α-$GP_1$) were designed for use in chronic efficacy studies in transgenic mice. Recombinant antibodies were transiently expressed in CHO cells, purified using standard Protein A affinity chromatography and desalted using PBS buffer. Endotoxin levels were confirmed to be < 10 EU/ml.

*Synthetic DPR*

Synthesis and purification of dipeptide repeat peptides was performed by Schafer-N (Copenhagen, Denmark): Sequences of the 15-repeat peptides were: $GA_{15}$: H-CHHHHHH$(GA)_{15}$-OH: $GP_{15}$: H-C$(GP)_{15}$-OH; $GR_{15}$: H-C$(GR)_{15}$-OH; $PA_{15}$: H-C$(PA)_{15}$-OH; $PR_{15}$: H-C$(PR)_{15}$-OH. Upon delivery from the supplier, the lyophilized peptides were dissolved in ≥ 99.9 % DMSO (Sigma) and stored at -20 °C. Five synthetic DPR peptides were used: $GA_{15}$ (2866 Da), $GP_{15}$ (2434 Da), $GR_{15}$ (3319 Da), $PA_{15}$ (2644 Da) and $PR_{15}$ (3921 Da).

*Indirect ELISA and $EC_{50}$ determination*

96-well-format microplates (Corning Incorporated, Corning, USA) were coated with DPR peptides at 5 or 20 µg/ml in coating buffer (15 mM $Na_2CO_3$, 35 mM $NaHCO_3$, pH 9.4). Blocking of non-specific binding sites was performed for 1 hrs at RT with PBS containing 0.1 % Tween-20 and 2 % BSA (Sigma-Aldrich). Primary human anti-DPR protein antibodies were diluted to the indicated concentrations and incubated for 1 hrs at RT, followed by incubation with an HRP conjugated donkey α-human IgG Fcγ-specific antibody (Jackson ImmunoResearch Laboratories Inc, West Grove, USA). Binding was determined by measurement of HRP activity in a standard colorimetric assay. $EC_{50}$ values were calculated by non-linear regression using GraphPad Prism (San Diego, USA).

*Kinetics using bio-layer interferometry*

Bio-layer interferometry (BLI) experiments were performed on an Octet RED96 instrument (Pall ForteBio LLC). Amine-reactive (AR2G) biosensors were used for covalent immobilization of $GA_{15}$, $GR_{15}$, $PR_{15}$ and $PA_{15}$ DPR peptides. AR2G biosensors were activated with EDC (1-Ethyl-3-[3-dimethylaminopropyl] carbodiimide hydrochloride; 20 mM in water; Pall ForteBio LLC) and s-NHS (N-hydroxysulfosuccinimide; 10 mM in water; Pall ForteBio LLC) for 300 s. This was followed by loading of the biosensor surface with 5 µg/ml peptides in 10 mM acetate buffer pH 6 (Pall ForteBio LLC) for 600 sec. After peptide loading, AR2G biosensors were quenched with 1 M ethanolamine pH 8.5 (Pall ForteBio LLC) for 300 s, rinsed with kinetics buffer (1:10 in $dH_2O$, Pall ForteBio LLC) for 120 sec (baseline) and antibody association was assessed at different concentrations in diluted kinetics buffer for 600 sec.

Antibody dissociation was evaluated in kinetics buffer for 600 sec. AR2G biosensors were regenerated for replicate experiments using 10 mM Glycine (Sigma) pH 2 and 10x PBS (Gibco) for neutralization. All binding data was referenced by collecting data with a kinetics buffer only reference. Unspecific binding to the sensor surface was tested using an isotype control antibody at the highest used concentration for each peptide. Streptavidin (SA) biosensors were used for immobilization of biotinylated $GP_{15}$. Biotinylation and purification of $GP_{15}$ was performed using a biotin labeling kit (Roche, Cat. no. 11 418 165 00) according to the manufacturer's protocol. For sensor loading, peptide-containing fractions from the purification column were pooled and diluted 1:3 in PBST. SA biosensors were activated using PBST before loading the biotinylated peptide for 600 sec. Upon peptide loading, SA biosensors were blocked for 300 sec with 0.1 % milk in PBST (assay buffer), rinsed in assay buffer for 180 sec (baseline) and antibody association was assessed at different concentrations in assay buffer for 600 sec. Antibody dissociation was evaluated assay buffer for 600 sec. SA biosensors were regenerated for replicate experiments using 10 mM Glycine pH 2 and 10x PBS. All binding data were referenced by collecting data with an assay buffer only reference. All experiments were performed at 25 °C. Data analysis was performed by using the Octet system software (Pall ForteBio LLC) with simultaneous $k_a/k_d$ global fitting with 1:1 interaction model. BLI sensorgrams were plotted with GraphPad Prism (San Diego, USA) upon fitting.

*Cell culture*

SH-SY5Y human neuroblast cell line (Sigma Cat. no. 94030304) was cultured in Ham's F12 (Sigma Cat. no. N488-500ML) and EMEM (Sigma Cat. No. M2279), 1:1 ratio, supplemented with 2 mM L-Glutamine (Gibco Cat. no. 25030-024), 1 % NEAA (Gibco Cat. no. 11140-035), 15 % heat-inactivated FBS (Gibco Cat. no. A31608-01, Gibco) and 1 % PS. Cell culture was maintained at 37 °C and 5 % CO2. U2OS human osteosarcoma cell line (ATCC Cat. no. HTB-

96) was cultured in McCoy's 5A (Sigma Cat. no. M4892) with 10 % heat-inactivated FBS (Gibco Cat. no. A31608-01) and 1 % Penicillin/Streptomycin. Cell culture was maintained at 37 °C and 5 % CO2. The motor neuron-like cell line NSC-34 (CELLutions Biosystems inc. Cat. no. CLU-140) was cultured in DMEM/F-12 no glutamine (Gibco Cat. no. 21331-020), supplemented with B27 (2X), 1X of N2, Glutamax (all supplements from ThermoFisher Sci.) and 10ng/ml of BDNF and GDNF (PeproTech) added freshly. Stable inducible T98G glioblastoma cell line (ATCC Cat. No. CRL-1690) expressing $GA_{161}$-GFP was cultured in EMEM (ATCC Cat. No. 30-2003) supplemented with 10% Tetracycline free FBS (Clontech Cat. No. 631367), 1X Antibiotic-Antimycotic (Gibco Cat. No. 15240062) and 2 μg/ml puromycin (Gibco Cat. No. A1113803). Expression of $GA_{161}$-GFP aggregates was induced by adding 2 μg/ml doxycycline (Sigma-Aldrich Cat. No. D9891) to the culture medium. Cell culture was maintained at 37 °C and 5% $CO_2$. Human neural stem cells (NSCs) were differentiated for 5-6 weeks resulting in functional neural networks (39). Shortly, NSCs were plated onto Matrigel-coated 6-well plates and grown in NSC media until reaching confluency. Media was then changed to D3 differentiation media and ~4-5 weeks upon induction of differentiation, differentiating neural cultures were sub-cultured onto Matrigel-coated 6-well plates for biochemistry, or 24-well μ-Plate (Ibidi Cat. no. 82406) for imaging. Using 0.05 % and 1X Defined Trypsin Inhibitor (Gibco Cat. no. R-007-100), cells were carefully re-suspended with 10 ml pipette to dissociate tissue-like clumps, centrifuged at 1000 RPM, 5 mins at room temperature (RT), re-suspended in D3 media and re-plated at ¼ ratio.

*$DPR_{50}$ plasmids*

In order to test the antibody DPR specificity via immunofluorescence, "DPR-only" genes were generated. DPR-only genes coding for 50 DPR repeats (47 for GP) were designed using all possible triplets to decrease the GC-content and avoid repetitions  and synthesized at GenScript flanked with NcoI and NsiI restriction sites, cloned in- and delivered in our deposited modified pUC18 Gateway entry clone plasmid (upstream of P2A-EGFP sequence) via NcoI and NsiI sites. DPR repeats were then C-terminally fused with EGFP via short GSG linker using Q5 site-directed mutagenesis kit (NEB Cat no. E0554S) and then cloned into our modified pSF-CAG-Kan (Oxford Genetics) Gateway destination vector termed pDEKA (Gateway cassette cloned from pEZY3 Addgene Cat. no. 18672 vector). GFP-only vector (generated via Q5 deletion of $GA_{50}$ from GA50-GFP) was used as a DPR-negative control. For experiments requiring DPR-only vectors without GFP, all 5 DPR sequences were PCR-amplified and restriction-cloned into pSME-CMV vector using BamHI and EcoRI sites (a generous gift from Paulo Paganetti) so that they contain N-terminal HA tag.

*NSC-34 transfection for the antibody screen*

To screen for the specificity of antibodies across each DPR, NSC-34 were cultured for 48 hrs in 24-well plates for imaging (IBIDI) pre-coated with 0.15 mg/ml Matrigel. 500 ng of plasmid and 0.83 µl Lipofectamine 2000 per well were used for transfection which was then performed according to the manufacturer information. Non-transfected cells and/or transfected with GFP-only as well as cells transfected with DPR but stained with secondary antibody only were present on each plate. 48 hrs post-transfection, cells were fixed for immunofluorescence analysis.

*Immunofluorescence (NSC-34 and neural culture)*

The medium was removed and cells were covered with 200 µl of 0.45 mg/ml Matrigel/medium mix for 30 mins at 37 °C, 5 % $CO_2$, to prevent cells to detach during immunofluorescence procedure. Cells were then washed once with PBS and fixed for 10-20 mins with 4 % paraformaldehyde in PBS. Cells were permeabilized and blocked during 1 hr at room temperature in PBS, 10 % donkey serum and 0.1% Triton X-100 (blocking/permeabilization buffer), then incubated with 25 nM of the tested human recombinant antibody at 4 °C overnight in blocking/permeabilization buffer. Cells were washed three times 10 mins in PBS 0.1 % Triton X-100 before incubation with α-human Alexa 594 and DAPI (0.1 µg/ml) in blocking/permeabilization buffer. Cells were washed 2 times in PBS 0.1 % Triton X-100 and once with PBS, then mounted with ProLong™ Gold Antifade Mountant with DAPI (Invitrogen Cat. no. P36931). Human neural cultures were fixed and stained as described in the NSC-34 immunofluorescence sub-section with rabbit α-Tuj1/ßIIITubulin (1:1000; Abcam Cat. no. ab18207), goat α-GFP-FITC (1:500; Rockland antibodies Cat. no. 600-102-215) primary antibodies. Donkey α-rabbit 546-Alexa fluor or Cy3-conjugated secondary antibodies (1:1000; Life Technologies Cat. no. A10040 and 1:1000, Jackson ImmunoResearch Cat. no. 711-165-152, respectively) were used to detect the rabbit primary antibodies and donkey α-mouse 647-Alexa fluor (1:1000 LifeTech Cat. no. A-31571) to detect the $^{ch}\alpha$-GA$_1$, $^{ch}\alpha$-GA$_3$ or an IgG murinized antibodies.

*SH-SY5Y Transfection and colocalization analysis*

To assess antibody uptake and target colocalization, adherent SH-SY5Y cells were cultured at an initial density of 70,000 cells per well in coated 24-well plates for imaging (Ibidi Cat. no. 82406, Ibidi). After 24 hrs cells were transfected with 1 µg (GA)$_{50}$-GFP or GFP control plasmids in 100 µl serum-free EMEM and 2 µl X-tremeGENE 9 DNA transfection reagent (Roche Cat. no. 25582900) per well, according to the manufacturer’s protocol. Six hrs after transfection, the culture medium was removed and replaced with medium containing 50 nM human antibodies α-GA$_1$, α-GA$_3$ or an IgG isotype control antibody. Cells were kept at 37 °C and 5 % CO2 and 72 hrs after antibody addition they were washed twice with PBS to remove non-internalized antibodies and fixed for 20 mins with 4 % paraformaldehyde. Fixed cells were permeabilized and blocked for 1 hrs with PBS containing 2 % BSA and 0.2 % Triton X-100. For antibody detection, cells were incubated with α-human Cy3 secondary antibody (Jackson Cat. no. 709-165-149) in PBS containing 2 % BSA for 1 hrs at room temperature. After washing twice with PBS, DAPI was added 1:1000 in PBS for 10 mins and, after a final PBS wash, cells were kept in PBS at 4 °C until imaging. Images were acquired on a Leica DMI6000 AFC Sp8 inverted confocal microscope at 10x magnification. Microscope settings used for acquisition were: 1024x1024 pixel image size, zoom = 1, speed = 400, Line/Frame average = 1. Sequential images were taken in three sequences using PMT sensors and the following lasers: DAPI - Diode 405, EGFP - Argon 488, Cy3 - DPSS 561. Images were analyzed with Imaris' "Coloc" algorithm (Bitplane, version 9.1.0) using a threshold of 40 for the red and green channel. Datasets from two independent experiments were combined for analysis. Each experiment was performed with two replicates wells per condition and three pictures at different positions from each well. Cropped pictures displayed in **Fig. 2A** were prepared with ImageJ (NIH, version 1.51 n) without intensity or contrast alterations from original images. Imaging was performed with equipment maintained by the Center for Microscopy and Image Analysis, University of Zurich.

**$GA_{161}$-GFP T98G glioblastoma antibody treatment, immunofluorescence and analysis:**

$GA_{161}$-GFP T98G cells were plated on laminin-coated German glass cover slips (Electron Microscopy Sciences Cat. No. 72298-02) in 24-well plate at a density of 50,000 cells/well consisting 2 μg/ml doxycycline in culture medium. Next day, cells were treated in triplicate wells with either α-$GA_1$ or IgG isotype control antibody at a final concentration of 100 nM for 72 h.

Following treatment, cells were washed three times with 1X HBSS containing calcium chloride and magnesium chloride (Gibco Cat. No. 14025-092) and fixed with 4% paraformaldehyde in

PBS at RT for 20-30 min. Cells were concurrently permeabilized and blocked in 1X DPBS, 0.2% Triton X-100, 2% BSA for 20 min at RT followed by three washes in 1X DPBS. Antibodies in the cells were stained with 1.5 µg/ml Alexa Fluor® 647-conjugated anti-human IgG (Jackson Immunoresearch Cat. No. 109-606-088) prepared in 1X DPBS, 2% BSA for 45 min at RT and washed three times with 1X DPBS. Finally, cells were treated with 1:10,000 Phalloidin-TRITC (Sigma Cat. No. P1951-.1MG) prepared in 1X DPBS, 2% BSA for 7-10 min at RT. Cells were washed three times with 1X DPBS, then mounted with ProLong Diamond AntiFade Mountant with DAPI (Invitrogen Cat. No. P36962).

Ten or eleven distinct 3D-fields spaced 0.27 µm apart were acquired for each sample with a 40X Oil-immersion lens (1.4 numerical aperture) with a Zeiss Axio Observer SD1 Inverted Microscope System (DAPI – 405 nm, $GA_{161}$-GFP – 488 nm, Phalloidin-TRITC – 561 nm and Alexa Fluor® 647 – 640 nm). Imaging system operation and image analysis was done using SlideBook 6 (Intelligent Imaging Innovations, USA). Cellular regions containing GA aggregates (GFP) or antibody (Alexa Fluor® 647) were masked by segmentation of the corresponding marker, setting a minimum intensity threshold and eliminating small objects (less than 10 voxels). Mask statistics were calculated for each field to assess the number and volume of GA aggregates and data was normalized to number of cells in the field.

*Flow cytometry (FACS)*

Human antibodies α-$GA_1$, α-$GA_3$ or an IgG isotype control were labeled with Alexa Fluor 488 (Molecular Probes #A10235) according to the manufacturer's protocol. SH-SY5Y cells were seeded in sterile 24-well cell culture plates (TPP) with 113,000 cells per well and kept at 37 °C and 5 % $CO_2$ for 24 h. The next day, cells were transfected with 1 µg $GA_{50}$-HA or incubated with 2 µl X-tremeGENE 9 DNA transfection reagent only (mock transfected). After another 24 hrs at 37 °C and 5 % $CO_2$, the culture medium was removed and labeled antibodies were added in fresh culture medium at 50 nM. Each $GA_{50}$-HA and mock-transfected condition was performed in triplicates. Cells were analyzed by flow cytometry 24 and 48hrs after antibody addition. Before measurements, cells were washed with PBS to remove remaining extracellular antibody and detached using 0.25 % Trypsin-EDTA (Gibco Cat. no. 25200-056). After resuspension in ice-cold medium cells were spun down at 450 *x* g and 5 °C. Supernatants were discarded, cell pellets resuspended in ice-cold PBS and transferred to a FACS tube through the cell strainer cap (Falcon Cat. no. 352235). To quench the fluorescence from membrane-bound antibodies Trypan Blue (Sigma Cat. no. T8154) was added to a final concentration of 0.04 % and the samples were put on ice. Flow cytometry analysis was performed on a FACSAria II (BD Biosciences) measuring 30,000 cells per sample. Cells were gated for viable and singlet cell populations and analyzed using Flowing Software 2 (Turku Centre for Biotechnology, version 2.5.1) to determine the number of Alexa 488 positive cells. Data plotting for illustrations and statistical analysis was performed using GraphPad Prism (GraphPad, version 7.03).

*Isolation of pure $GA_{50}$-GFP aggregates, scanning electron microscopy (SEM) imaging and correlative light-electron microscopy (CLEM)*

4 million of HEK239T cells were plated onto Matrigel-coated 10 cm culture dish in “OHN media” (Opti-MEM Gibco Cat. no. 11058-021 supplemented with B27- Cat. no. 12587-010 (0.5X), N2 Cat. no. 175020-01 (0.5X), Glutamax Cat. no. 35050038 (1X) and bFGF Cat. no. PHG0261 (all supplements from ThermoFisher Sci.)). 24 hrs later, cells were transfected with 15 µg $(GA)_{50}$-GFP plasmid using 21 µl Lipofectamine 2000 (ThermoFisher Sci Cat. no. 11668019) in OHN medium. Medium was exchanged 12 hrs post transfection. 72 hrs post transfection, spent medium was removed from cells and aggregates were extracted as follows: cells were lysed in the dish with 1 ml of 1 % sarkosyl HSI lysis buffer. Lysate was then

transferred into 15 ml falcons and incubated on ice for 10 mins with three rounds of mild 10 sec vortexing. Lysate was then brought up to 10 ml with 1 % sarkosyl HSI buffer and centrifuged at 300 *x* g for 10 mins at 4 °C. Supernatant was then carefully removed and pellet was re-suspended in 5 ml of HSI buffer, centrifuged at 300 *x* g for 5 mins at 4 °C. Supernatant was then carefully removed and the pellet was washed with 5 ml of $ddH_2O^{++}$ (Invitrogen Cat. no. 10977-035) supplemented with cOmplete EDTA free protease inhibitor cocktail (Roche Cat. no. 11873580001) and PhosSTOP (Roche Cat. no. 04906845001) (1 tablet and ½ tablet per 10 ml, respectively). The washed pellet was then centrifuged at 300 *x* g for 5 mins at 4 °C and finally re-suspended in 25 µl of $ddH_2O^{++}$. 15 µl of pure aggregates were then pipetted onto glow discharged glass coverslip (10 mm) and the water was left to evaporate at RT inside chemical hood ON. For antibody incubation experiments, cells were prepared as described above, however the cell extract was prepared in HSI buffer without detergent as follows: spent media was removed, cells were detached into 1 ml of HSI using cell scraper and the cell suspension was then transferred into 1.5 ml protein low-binding Eppendorf tubes (Cat. no. 7000228) and sonicated at 40 % amplitude for 2 mins of sonication time (5 s–on, 5 s–off pulses) in ice-cold water in water-bath sonicator (Cat. no. Q500A-110, QSonica). Resulting cell extracts were then treated with 2 µl Benzonase (Millipore Cat. no. E1014-5KU) and 2 mM MgCl2 and incubated with 0.2 µM of $\alpha$-$GA_1$ or IgG-control antibodies (+ non treated control) at 37 °C for 24 hrs while shaking at 150 r.p.m. (Thermomixer, Eppendorf). Cell extract was then diluted with 2 % sarkosyl HSI and pure $GA_{50}$-GFP aggregates were then isolated and prepared for IF and subsequent SEM imaging by pipetting onto a coverslip as described above. Immunofluorescence (IF; GFP and far-red channels) and brightfield imaging were done using IN Cell Analyzer 2500HS widefield microscope imaging the entire coverslips with 40X air objective. Samples were sputter coated with 3 nm of platinum (Safematic, Bad Ragaz, Switzerland Cat. no. CCU-010). SEM imaging was done at the Center for Microscopy and

Image Analysis UZH using an Apreo VS SEM microscope (Thermofisher Scientific, Eindhoven, The Netherlands), and imaged at an acceleration voltage of 4 kV at a pixel size of 10nm using an Everhardt Thornely secondary electron detector and the Maps software package. SEM data were analyzed using Maps viewer (Thermo Fisher) and CLEM was performed (**Fig. S15*E***) using both Maps viewer and offline version of INCell Analyzer (GE Healthcare) software to visually confirm GFP and / or AF647 positivity of the high-resolution SEM pictures of $GA_{50}$-GFP aggregates. Only aggregates that were $GFP^+$ and $AF647^+$ (antibody-treated conditions) were further processed. Each aggregate (NT, n=139; IgG, n=125; α-$GA_1$, n=183; α-$GA_3$, n=193) was manually traced using a graphic tablet and the total occupied area was measured using ImageJ. Quantification of the surface of the antibody treated $GA_{50}$-GFP aggregates was done by assessing the porosity of the aggregates using trained ilastik algorithm to segment pixels followed by pixel measurements of thresholded images in ImageJ. Statistical analysis was then performed in Prism (GraphPad) and one-way ANOVA followed by Tukey's multiple comparison test was applied on the datasets.

*Antibody uptake and $GA_{50}$-GFP aggregation in human neural culture*

Human neurons were differentiated as described above. Cells were transduced between day 22-34 of differentiation with an all-in-one Tet-ON lentiviral vector (LV) conditionally expressing 50 repeats of GA, C-terminally fused with a GS-linker to the enhanced Green Fluorescent Protein (vLVX-$GA_{50}$-GFP; $GA_{50}$-GFP generated as described above). $GA_{50}$-GFP is under the control of Tet-responsive promoter $P_{tight}$, which consists of seven tet operator sequences followed by the minimal CMV promoter, ensuring its tight control by doxycycline (DOX). vLVX-$GA_{50}$-GFP further contains a red fluorescent protein mRuby2, linked via self-cleaving P2A peptide sequence to the improved tetracycline-controlled transactivator (rtTA) under the control of a core of the constitutive EF-1α promoter. Faint mRuby2 signal can be used to visualize live transduced cells while in culture and is bleached rapidly post-PFA fixation which allows for use of the red channel for immunofluorescence staining. Transduced human neural cultures were sub-cultured 5 days post-transduction (at 1/3-¼ ratio into 24-well Ibidi imaging plates for IF or 6-well plates for biochemistry). Antibody treatment with 0.2 μM of $^{ch}\alpha$-GA$_1$, $^{ch}\alpha$-GA$_3$ or IgG control antibodies, as well as the induction of expression of GA$_{50}$-GFP with 1 μg/ml of Doxycycline (DOX; Clontech Cat. no. 631311), was initiated 6-7 days post-subculture via addition into the culture media until the endpoint except for the 2 weeks ON/ 1 week OFF condition where DOX was removed for the last week of culture while the treatment antibodies were kept. Media was changed every 3 days. At 3, 7 and 21 days, cells were fixed and stained as described in the Immunofluorescence (*NSC-34 and neural culture)* subsection. For determining the localization of antibodies in the different intracellular compartments (lysosomes and late endosomes), neurons were treated for 3 days with 0.2μM chimeric antibodies. Neural cultures were imaged on Nikon Eclipse Ti-E confocal, Leica SP5, Leica SP8 inverted confocal or GE InCell Analyzer 2500 HS widefield microscopes.

*Distance analysis for GA$_{50}$-GFP colocalization with intracellular vesicles.*

GA$_{50}$-GFP expressing neural cultures cultured on coverslips (thickness #1.5H) were fixed for 20 mins in 4 % PFA and stained with α-GFP-FITC (Abcam Cat. no. ab290) and either α-RAB7 (Cell Signaling Cat. no. 2094S) or α-LAMP1 (Abcam Cat. no. ab21470) (1 in 400) and detected with α-Rabbit 594. Cells were imaged on a Leica Sp8 Inverse STED at 93x. The 592 and 775 nm wavelength STED lasers were used to deplete the green and red channels respectively. 15-17 regions of interest with comparable number of GA$_{50}$-GFP particles were analyzed per condition ($^{ch}\alpha$-GA$_3$ or IgG) for LAMP1 and RAB7 using the "surface creation" and the incorporated Matlab extension "distance transformation" in Imaris (Bitplane). Measured distances are plotted and analyzed using GraphPad Prism7.

*Viability assay*

Human neural cultures expressing an inducible $GA_{50}$-GFP construct were cultured in a flat-bottom white 96-well plate. $GA_{50}$-GFP expression was induced with 1 µg/ml of DOX (Clontech Cat. no. 631311) and at the same time, cells were treated with 0.2 µM of $^{ch}\alpha$-$GA_{1,3}$ or with the corresponding IgG isotype control for 3 days. Non-induced control was treated with $^{ch}\alpha$-$GA_1$ antibody. To perform the viability assay, the whole medium was removed from the well, and immediately replaced by 50 µl of warm D3 neural medium. Negative control was performed in wells containing only medium. Reagents from the Real Time-Glo™ MT Cell viability assay (Promega Cat. no. G9711) were diluted in D3 medium according to manufacturer's recommendations and added to the wells reaching final volume of 100 µl per well. Cells were kept at 37 ºC, 4% $CO_2$ and luminescence was measured 60 minutes later on a pre-heated Tecan Infinite200 device.

*GA-insolubility biochemistry assay – human neural culture and AAV($G_4C_2$) mouse brains (SarkoSpin)*

Cells were collected by scraping in 100 µl cell lysis buffer (0.5 % sarkosyl (Sigma Cat. no. L5125-100G) + 0.5 µl Benzonase (Millipore Cat. no. E1014-5KU) and 2 mM MgCl2 in HSI buffer (10 ml of HSI: 10 mM TRIS, 150 mM NaCl, 0.5 mM EDTA, 1 mM DTT + 1 tablet of cOmplete EDTA free protease inhibitor cocktail (Roche Cat. no. 11873580001) and ½ tablet of PhosSTOP (Roche Cat. no. 04906845001). Re-suspended samples were transferred into protein low binding Eppendorf tubes. The wells were then washed with additional 100 µl of lysis buffer and transferred to the corresponding tube, to collect the remaining cells. From the obtained total cell lysate, 30 µl were used for total blots and the remaining sample was further processed to obtain supernatant and aggregate-containing fractions. To 170 µl of the remaining cell homogenate, 178 µl HSI with 4 % sarkosyl and 52 µl HSI was added (to obtain a final sarkosyl concentration of 2 % in total 400 µl volume). Similarly, the entire frozen right hemispheres of AAV($G_4C_2$) mice were homogenized at 20 % w/v in HSI via three rounds of 30 seconds homogenization in tubes containing a mixture of ceramic beads using a Minilys device (Bertin) at full speed with cooling on ice between each step. Samples were aliquoted (150 µl), snap frozen in liquid nitrogen and kept in -80ºC before use. Brain homogenates were then thawed on ice and placed in 4 % sarkosyl lysis buffer (similarly as above) to obtain 2 % sarkosyl in 400 µl final volume. The samples were incubated at 37 °C, 600 rpm for 45 mins (Thermomixer, Eppendorf), and after increasing the volume by addition of 200 µl HSI, the samples were vortex mixed and centrifuged at 21,200 *x* g on a benchtop centrifuge (Eppendorf) for 30 mins at room temperature. Supernatants (~450 µl) were transferred to a new tube and the remaining supernatant was carefully removed to leave the pellets completely dry. The latter were then re-suspend in 80 µl (cells) or 125 µl (brains) of 0.5 % sarkosyl HSI and analyzed by filter retardation assay or immunoblot as described below.

*SDS-PAGE and Immunoblotting*

For total cell lysates and lysates of SNS pellets, protein concentration was adjusted using Pierce™ 660 nm Protein Assay Reagent (Thermo Fisher Scientific Cat. no. 22660) and 20 µg of total protein per well was used for immunoblots. Samples were re-suspended in final 1X LDS loading buffer (LifeTech Cat. no. NP0007) with 1X final Bolt sample reducing agent (Cat. no. B0009), denatured at 95ºC for 10 mins, and loaded on Bolt 4-12 % or 12 % Bis-Tris (Cat. nos. NW04122BOX, NW00122BOX, NW00125BOX) for SNS samples. For immunoblots, gels were transferred onto nitrocellulose membranes using iBlot® 2 Transfer NC Stacks (Cat. no. IB23001) with iBlot 2 Dry Blotting System (Cat. no. IB21001). To assess sarkosyl-insoluble $GA_{50}$-GFP fractions, re-suspended pellets were loaded onto two 0.1 µm nitrocellulose membranes in a filter retardation device (dot blot). Vacuum was applied to the device to force samples to pass through the membranes. Membranes were then blocked with 5 % w/v non-fat skimmed powder milk in 0.05 % v/v Tween-20 (Sigma Cat. no. P1379) in PBS (milk PBST) and probed with primary antibodies (rabbit α-GFP antibody (1:5000; Abcam cat. no. ab290), mouse α-actin (1:5000; Sigma Cat. no. A5441), rabbit α-UBK48 (1:1000; Millipore cat. no. 05-1307), rabbit anti-HA (1:10000; Proteintech Cat. no. 51064-2-AP), rabbit α-poly-GR (1:1000; generated in house RB7635)) overnight in PBST with 1 % w/v milk, washed three times with PBST, followed by incubation with secondary HRP-conjugated goat anti mouse or rabbit AffiniPure IgG antibodies (1:5000, 1:10000, respectively) (Jackson Immuno Research Cat. nos. 115-035-146 and 111-035-144, respectively) in 1 % milk PBST. After three washes in PBST, immunoreactivity was visualized by chemiluminescence using SuperSignal West Pico or Femto Chemiluminescent Substrate (PierceNet Cat. nos. 34077, 34096) on Amersham Imager 600RGB (GE Healthcare Life Sciences Cat. no. 29083467). Fiji was used to quantify the dot and western blot signal intensity. For dot blot, intensity values were normalized to the average intensity value of non-treated samples (3-day time point).

*Image analysis for quantification of $GA_{50}$-GFP structures*

20x widefield images obtained from the GE InCell Analyzer 2500 HS widefield microscope were analyzed for quantification of extracellular and intracellular $GA_{50}$-GFP. 36 images per well, equally spaced and centered on the middle of the well were acquired (3 to 6 wells per condition). Nuclei count was performed by thresholding the image, dilating the binary mask and particles were automatically counted by using Fiji. $GA_{50}$-GFP extracellular structures were manually counted on 5 images per well, in 3 wells per condition and normalized to the number of nuclei. Since extracellular GA immune complexes were confused with intracellular $GA_{50}$-GFP structures when using a simple intensity threshold method for quantification (data not shown), we trained Ilastik (version 1.3.0), a supervised machine learning software, to identify intracellular $GA_{50}$-GFP structures. Ilastik was trained in two separate analyses on a subset of images until it correctly segments bright intracellular aggregates in one analysis and intracellular inclusions in an independent analysis. The pixel classifications were then extended to the whole set of GFP channel images (108-216 images per condition). As an output, Ilastik created two distinct binary masks per image, that were automatically counted using Fiji. Intracellular structures (bright aggregates and inclusions) (**Fig. 4*C***) were normalized to the number of nuclei, summed and represented as intracellular $GA_{50}$-GFP structures.

*C9ORF72 transgenic mice*

The Massachusetts General Hospital Institutional Animal Care and Use Committee (IACUC) approved animal care and experimental protocols. A cohort of 200 transgenic mice expressing 450 *C9ORF72* $G_4C_2$ repeats (referred to $C9^{450}$) (37) was generated at Charles River by *in vitro* fertilization and 50 Wild type C57BL6/J mice age-matched were purchased from Charles River. Mice were aged at the Harvard Biological Research Infrastructure for immunotherapy studies. $C9^{450}$ mice accumulate age-dependent DPR pathology and mild behavior deficits. At three months old, 160 mice were randomized in five weight and gender randomized groups for chronic efficacy studies, referred as Wt PBS (n = 40), $C9^{450}$ PBS (n = 40), $C9^{450}$ α-$GA_1$ (n = 40), $C9^{450}$ α-$GA_3$ (n = 40) and $C9^{450}$ α-$GP_1$ (n = 40). These mice received weekly 30 mg/kg of chimeric DPR antibodies intraperitoneally in phosphate buffer saline 1x for 4 or 16 months. Half of the animals were euthanized at 7 months old (4 months treatment) and brains were collected for RNA and DPR quantifications by reverse transcription- quantitative PCR (rt-qPCR) and immunoassays, respectively. The remaining animals went through behavioral testing at 13 and 18 months of age and were subsequently sacrificed for brain biochemistry and histology analyses. Male and female mice were tested for marble burying and open field. A significant deficit was observed only for open field in $C9^{450}$ males compared to control littermates (**Fig. 7*B***, *SI Appendix,* **Fig. S15*F*** ). All the mice were bled every 8 weeks for serum antibody titers along the study. A subset of $C9^{450}$ and Wt mice were divided into five groups (Wt-PBS (n = 3), $C9^{450}$ Isotype (n=3), $C9^{450}$ α-$GA_1$ (n = 3), $C9^{450}$ α-$GA_3$ (n = 3) and $C9^{450}$ α-$GP_1$ (n = 3)) and aged until 10 months old for pharmacokinetics studies. A single intraperitoneal injection of 30 mg/kg was performed per mouse before plasma and brains were collected at different time points: -7 d, 0 (injection time point), +1 h, +8 h, 24 h, +2 d, +4 d, +7 d, +10 d, +14 d and +21 d for plasma and +2 d, +7 d and +21 d for brains ("d" referred for days and "h" for hours). An additional cohort of 20-month-old mice exhibiting high level of DPR pathology were used for target engagement with collection of brains 10 days after a single intraperitoneal injection of IgG isotype control, α-$GA_1$, α-$GA_3$ and α-$GP_1$ (*SI Appendix,* **Fig. S1*E***).

*Neonatal AAV($G_4C_2$)$_2$ and AAV($G_4C_2$)$_{149}$ viral injections*

Generation of AAV($G_4C_2$)$_2$ and AAV($G_4C_2$)$_{149}$ and intracerebroventricular (ICV) injections of these AAV were done as previously described in (38). Briefly, for AAV generations, we inserted the ($G_4C_2$)$_2$ and ($G_4C_2$)$_{149}$ repeats with 119 base pairs of the 5' flanking region and 100 base pairs of 3' flanking region of the *C9ORF72* gene into the HindIII and XhoI restriction sites of the adeno-associated virus (AAV) expression vector pAM/CBA-pl-WPRE-BGH containing inverted repeats of serotype 2. AAV($G_4C_2$)$_2$ and AAV($G_4C_2$)$_{149}$ particles were packaged into serotype 9 type capsid and purified using standard methods as in (38). The genomic titer of each virus was determined by qPCR. The solutions of AAV were diluted with sterile PBS 1x. Briefly, for AAV ICV injections, a 32-gauge needle (Hamilton Company) attached to a 10 μl syringe (Hamilton Company) was inserted at approximately two-fifths the distance between lambda suture and each eye of C57BL/6J pups at post-natal day 0 after they were cryo-anesthetized on ice until pups exhibited no movement. Two microliters (1E10 genomes/µl) of AAV($G_4C_2$)$_2$ or AAV($G_4C_2$)$_{149}$ solution was manually injected into each cerebral ventricle. After injections, pups were placed on a heating pad until they completely recovered from anesthesia and then were placed back into their home cages. In total, we generated a cohort of 90 mice injected with AAV($G_4C_2$)$_{149}$ and 50 mice injected with AAV($G_4C_2$)$_2$ for efficacy studies testing α-$GA_1$. At two months old, four different groups were randomized by gender and weight (referred as ($G_4C_2$)$_2$ isotype, ($G_4C_2$)$_{149}$ isotype, ($G_4C_2$)$_2$ α-$GA_1$ and ($G_4C_2$)$_{149}$ α-$GA_1$) and they were intraperitoneally injected weekly with 30 $mg.kg^{-1}$ of α-$GA_1$ or isotype control for two or ten months. Four and 12-month-old animals with chronic administration of antibodies were euthanized for DPR immunoassays and histology.

*Open field assay*

$C9^{450}$ or AAV($G_4C_2$) male and female mice were placed in the center of an open field area (Perspex box of 11 1/8'', WxLxH) and were allowed to explore the area for 15 mins. The movement was monitored through the use of an overhead Noldus camera with Ethovision XT software. Mice were tracked for multiple parameters, including total distance traveled, average speed and time mobile. Data were analyzed with Prism GraphPad 6.0 (GraphPad Software) using nonparametric Kruskal-Wallis test compare with the mean rank of Wt group injected with PBS (*$P < 0.05$, ** $P < 0.001$, *** $P < 0.0001$)

*Inverted grid assay*

AAV($G_4C_2$)$_2$ and AAV($G_4C_2$)$_{149}$ female mice were analyzed by inverted grid at 9 months of age. Male animals were excluded from the analysis since they were not able to perform the test and fell from the grid within a few seconds independently of their genotype. Mice were brought into the examination room and allowed to acclimate for 15 minutes before the test was started.

The inverted grid test uses a metallic wire grid to measure muscular strength and conditioning over time, as assessed by the sustained ability to remain hanging from the grid. The test measures the hang time of the 4 limbs in seconds as well as the "holding impulse" (Holding Impulse = Body Mass x (Gravity) 0.00980665 N/g x Hang Time) for normalization to body weight. Each mouse was placed on the grid for 5 seconds to adjust. The grid was then inverted and held approximately 45 cm over a mouse cage containing 6 cm of bedding for a maximum time of 5 minutes. Each holding period begins with all four paws of the mouse firmly grasping the grid. The hang time was defined as the amount of time that elapses before the mouse falls down from the inverted grid and was measured visually with a stopwatch. Each session, the procedure was repeated three times with a maximum rest time of 5 mins between each assessment. All mice were assessed over 3 days, with 3 procedures per day, to evaluate baseline capabilities and learning.

*Histology*

Human-derived antibodies against DPRs were tested at 1 and 25 nM on PFA 4 % fixed paraffin mice sections (8 µm, 20 months old) and 8 µm paraffin sections from cerebellum of C9ORF72 patients or healthy individuals (**Fig. 1** and **Table 1**). HRP conjugated human secondary antibody (Jackson) was used at 1/175 dilution.

Treated $C9^{450}$ and $AAV(G_4C_2)_{149}$ mice were euthanized by CO2 asphyxiation and perfused with cold PBS 1x before tissue collections. For efficacy studies, the brain was removed and processed as follows: the left hemisphere was snap-frozen and stored at -80 °C for q-PCR *C9ORF72* RNA and DPR quantification by immunoassays. The right hemisphere was post-fixed in paraformaldehyde 4 % for 24 hrs and embedded in paraffin for sagittal immunostaining (10 µm) following protocols described in (37) with in-house generated antibodies against poly-GA (Cat. no. Rb4334, 1:1000; or human $\alpha$-$GA_1$ for co-immunostaining in *SI Appendix*, **Fig.**

**S12*A*** and ***B***); poly-GP (Cat. no. Rb4335,1:1000), poly-GR (Cat. no. Rb4995,1:1000), N-term-poly-GA (Cat. no. RB9261, 1:122; raised against the peptide MELRSRAL translated in the poly-GA frame) and rabbit immunodetection kit (Vector Laboratories).

Representative images after immunofluorescence staining were captured using a Nikon Eclipse TI confocal microscope and the NIS Elements AR software. The images were taken as 3-channel RGB color (TRITC = red, FITC = green, DAPI = blue). Four images (40x) were acquired for each sagittal brain section, two of the CA1 hippocampus and two of the motor cortex. Images for poly-GA and poly-GP were quantified with a custom high-throughput MATLAB script (https://github.com/krsavage/Confocal-image-analysis-for-protein-aggregates-GFAP-and-IBA1). The program separates each image into the FITC/green channel, converts it to grayscale, then background-subtracts it using morphological Tophat filtering. This creates a binary image with clusters, which are analyzed as connected components by count, size, and percent area ((true_pixels/total_pixels)*100)).

Phospho-TDP-43 immunohistochemistry was performed as described in (42).

*RNA extraction and quantitative RT-PCR*

Total RNA from mouse brain was isolated with TRIZOL (Invitrogen) and first-strand complementary DNA (cDNA) was synthesized using the SuperScript III First-strand synthesis kit (Thermofisher). Quantitative RT-PCR reactions were conducted and analyzed on a CFX96 touch Real-time PCR machine (Bio-Rad). Human *C9ORF72* transgene expression levels were determined using the SYBR Green Supermix (Bio-Rad). Total human C9ORF72 were determined using TaqMan real time-PCR and normalized to glyceraldehyde-3-phosphate dehydrogenase (GAPDH). Primers and probe sequences for the Human total C9ORF72 are: Forward TGTGACAGTTGGAATGCAGTGA, reverse GCCACTTAAAGCAATCTCTGTCTTG and probe TCGACTCTTTGCCCACCGCCA and for mouse GAPDH mRNA are: Forward GGCAAATTCAACGGCACAGT, reverse GGGTCTCGCTCCTGGAAGAT and probe AAGGCCGAGAATGGGAAGCTTGTCATC.

*Serum and brains antibody titers*

96-well microplates (Corning Incorporated, Corning, USA were coated with either synthetic $GA_{15}$ or $GP_{15}$ DPR peptides at 0.5 µg/ml (for serum drug levels) or 1 µg/ml (for brain drug levels) in coating buffer (15 mM $Na_2CO_3$, 35 mM $NaHCO_3$, pH 9.4) overnight at 4 °C. Blocking of non-specific binding sites was performed for 1 hr at room temperature with PBS containing 0.1 % Tween-20 and 2 % BSA (Sigma-Aldrich). Serum samples diluted to 1:40,000 in PBS or 30 µg total protein from brain homogenates were added and incubated for 2 hrs at room temperature, followed by incubation with an HRP-conjugated donkey α-human IgG1-specific antibody (Jackson, 1:10,000). Antibody standard curves were prepared by two-fold serial dilutions of α-$GA_1$, α-$GA_3$, or α-$GP_1$ in PBS/mouse serum with an initial antibody concentration of 1 nM. Binding was determined by measurement of HRP activity in a standard colorimetric assay. Estimated drug serum and brain levels were determined by linear regression using Excel 2016 (Microsoft, USA).

*Poly-GA, poly-GP and poly-GR immunoassays* (*SI Appendix,* **Fig. S11** and **Fig. S13*G***)

Mouse brains were thawed on ice in approximately 5 x (w/v) lysis buffer (150 mM NaCl, 20 mM Tris, pH 7.5, 1 mM EDTA, 1 mM EGTA, 1 % Triton X-100, 0.2 % sodium deoxycholate) with protease inhibitor cocktail (cOmplete, Sigma). Homogenization was performed using MP BIO FastPrep-24 with Lysing matrix D 1.4 mm ceramic beads (MP Biomedicals, Solon, OH, USA) according to manufacturer's protocol. Homogenates were sonicated for 10 seconds in 4 °C water bath and then separated from beads and debris removed by centrifugation at 850 *x g* for 10 mins at 4 °C. The homogenate was clarified by centrifugation at 16,000 *x g* for 20 mins at 4 °C. Total protein concentration was determined using BCA protein assay (Thermo Scientific) and used for normalization of poly-GA and poly-GP levels. The lysate was further fractionated by ultracentrifugation at 100,000 *x g* for 30 mins at 4 °C. The pellet was resuspended in 2 % SDS and heated to 95 °C for 5 mins.

Poly-GA, poly-GP and poly-GR levels were measured as previously published (12). Poly-GA levels in lysates were measured with an MSD-based poly-GA sandwich immunoassay that employed an in house-generated rabbit polyclonal poly-GA antibody (Rb4333) and a mouse monoclonal poly-GA antibody (clone 5F2). MSD-based sandwich immunoassays for poly-GP or poly-GR utilized affinity-purified rabbit polyclonal antibodies for each DPR as capture and detection antibodies. Response values corresponding to the intensity of emitted light upon electrochemical stimulation of the assay plates using the MSD QUICKPLEX SQ120 were acquired and background corrected using the average response from lysates obtained from non-transgenic mice.

*Poly-GA immunoassay with denaturation of samples* (**Fig. 5**)

Poly-GA in mouse brain lysates was measured using a Meso Scale Discovery sandwich immunoassay. In this assay, $^{ch}\alpha$-GA$_3$ was used as capture antibody, and human α-GA antibody α-GA$_4$ along with a SULFO-tag α-human secondary antibody was used for detection. Prior to measurements, SDS and TCEP were added to samples to final concentration of 1 % and 1 mM, respectively, and heated to 95 °C for 5 mins to dissociate treatment antibody from poly-GA. This treatment was shown to eliminate interference from antibody-GA complexes. Poly-GA concentrations were interpolated from the standard curve using 60X-GA expressed in HEK 293 cells and expressed as ng/mg total protein.

**Supplementary Figure Legends**

**Table S1. Summary of the characterization of human-derived antibodies against DPRs**

Affinity, kinetics and specificity of human-derived monoclonal anti-DPR antibodies determined by enzyme-linked immunosorbent assay (ELISA), biolayer interferometry (BLI), immunohistochemistry (IHC) and immunofluorescence (IF). Kinetic data is shown for the primary DPR target as determined by ELISA and indicated with an asterisk (*). $K_D$ equilibrium dissociation constant, $k_a$ association rate constant and $k_d$ dissociation rate constant.

**Fig. S1. Specificity and binding kinetics of human-derived antibodies against DPRs**

**(A)** Indirect enzyme-linked immunosorbent assay (ELISA) binding curves of human-derived antibodies to synthetic 15-mers of GA, GP, GR, PA, PR or bovine serum albumin (BSA) shown as the mean of duplicates. **(B)** Kinetic binding curves generated by biolayer interferometry. Curves display 600 sec of antibody binding to 15-mers of GA, GP, GR, PA or PR followed by a 600 sec dissociation phase. Fitting curves from a 1:1 binding model are shown with red solid lines for each concentration. All graphs are presented as 1:1 dilution series (from highest to lowest concentration: blue, purple, green, red and orange) of antibody with the highest concentrations, depicted in a solid blue line being: $\alpha$-GA$_1$ = 30 nM, $\alpha$-GA$_2$ = 30 nM, $\alpha$-GA$_3$ = 30 nM, $\alpha$-GA$_4$ = 20 nM, $\alpha$-GP$_1$ = 20 nM, $\alpha$-GR$_1$ = 20 nM, $\alpha$-PR$_1$ = 20 nM, $\alpha$-PR$_2$ = 20 nM, $\alpha$-PR$_3$ = 20 nM, $\alpha$-PA$_1$ = 30 nM and $\alpha$-PA$_1$ = 30 nM. Dashed black lines indicate data from a duplicate experiment for the highest antibody concentrations in each graph.

**Fig. S2. Characterization $\alpha$-GA$_1$ and $\alpha$-GA$_3$ antibodies binding specificity in tissues and motor-neuron-like cells**

**(A, B)** Immunohistochemistry on 8 μm sections of post-mortem cerebellum molecular layer from a C9ORF72 ALS/FTD patient and healthy control (upper panels) or the caudate putamen of C9$^{450}$ mice and wild-type littermate (lower panels) using the human-derived antibodies α-

$GA_1$ **(A)** and α-$GA_3$ **(B)** detected with a secondary α-human IgG antibody coupled to the enzyme horseradish peroxidase (HRP). Nuclei were stained with hematoxylin. Scale bars = 10 μm and 5 μm (insets). **(C, D)** Confocal images of NSC-34 motor neuron-like cells 48 hrs after transfection with either one DPR species (50 repeats) tagged to GFP or with GFP-only. **(C)** Cells were fixed and stained with DAPI and 25 nM of the human antibody α-$GA_1$, or **(D)** 25 nM of the human antibody α-$GA_3$ detected with α-human IgG antibody coupled to Alexa 594 (red). Scale bars = 20 μm.

**Fig. S3. Characterization of α-GA binding specificity in motor-neuron like cells**

Confocal images of NSC-34 48 hrs after transfection with one DPR species (50 repeats) tagged to GFP. **(A-B)** Cells were fixed and stained with DAPI and 25 nM of the human antibody α-$GA_2$ **(A)**, or 25 nM of the human antibody α-$GA_4$ **(B)** detected with α-human IgG antibody coupled to Alexa 594 (red). Scale bars = 20 μm.

**Fig. S4. Characterization of α-GP binding specificity in motor-neuron like cells**

Confocal images of NSC-34 48 hrs after transfection with either one DPR species (50 repeats) tagged to GFP or with GFP-only or non-transfected (NT). **(A)** Cells were fixed and stained with DAPI and only the secondary α-human IgG antibody coupled to Alexa 594 (red). **(B)** Cells were stained with DAPI and 25 nM of the human antibody α-$GP_1$ detected with α-human IgG antibody coupled to Alexa 594 (red). Scale bars = 20 μm.

**Fig. S5. Characterization of α-PA binding specificity in motor-neuron like cells**

Confocal images of NSC-34 48 hrs after transfection with one DPR species (50 repeats) tagged to GFP. **(A-B)** Cells were fixed and stained with DAPI and 25 nM of the human antibody α-$PA_1$ **(A)**, or 25 nM of the human antibody α-$PA_2$ **(B)** detected with α-human IgG antibody coupled to Alexa 594 (red). Scale bars = 20 μm.

**Fig. S6. Characterization of α-PR binding specificity**

Confocal images of NSC-34 48 hrs after transfection with one DPR species (50 repeats) tagged to GFP. **(A-B)** Cells were fixed and stained with DAPI and 25 nM of the human antibody α-PR$_2$ **(A)**, or 25 nM of the human antibody α-PR$_3$ **(B)** detected with α-human IgG antibody coupled to Alexa 594 (red). Scale bars = 20 µm.

**Fig. S7. Intracellular expression of GA$_{50}$ increases uptake/retention of poly-GA specific antibodies in SH-SY5Y cells**

**(A)** Confocal fluorescence images of cultured SH-SY5Y human neuroblastoma cells transfected with a GFP control plasmid and incubated with 50 nM of human α-GA$_1$, α-GA$_3$ or an IgG isotype control antibody for 72 hrs. Antibodies were visualized after fixation with a secondary α-human IgG antibody (red) and nuclei were stained with DAPI (blue). Scale bar = 100 µm. **(B)** Image quantification of the percentage of area covered by antibody that colocalized with GFP. Each data point represents the mean value of one well calculated from three separate pictures. N = 4 biological replicates from 2 independent experiments. **(C)** Quantification of flow cytometry analysis of HA-GA$_{50}$ or mock transfected SH-SY5Y cells incubated with Alexa 488-labeled α-GA$_1$, α-GA$_3$ or IgG isotype control for 24 hrs. Each data point represents 30,000 cells and the experiment was performed with biological triplicates (n = 3). **(D)** Flow cytometry scatter plots showing non-transfected cells treated with Alexa 488-labeled α-GA$_1$, α-GA$_3$ or an IgG isotype control after 48 hrs. Dotted line indicates the cutoff between Alexa 488-positive and negative cells (as determined in Fig 2D). Each plot represents 30,000 analyzed cells. **(E)** Quantification of flow cytometry analysis of untransfected SH-SY5Y cells incubated with Alexa488-labeled α-GA$_1$, α-GA$_3$, IgG isotype control or medium only for 24 or 48 hrs. Each data point represents 30,000 cells and the experiment was performed with biological duplicates (n = 2). **(B, C)** Horizontal lines represent the mean ± SD, one-way

ANOVA followed by Tukey's multiple comparison test. No indication $P > 0.05$, * $P \leq 0.05$, ** $P \leq 0.01$, *** $P \leq 0.001$.

**Fig. S8. Antibody internalization and colocalization with $GA_{50}$-GFP in human neuronal culture.**

**(A)** Confocal immunofluorescence images of a 6-week-old human neuronal culture non-transduced and treated for 3 days with 0.2 µM of $^{ch}\alpha$-$GA_3$ antibody (upper panel) or non-treated (lower panel). $^{ch}\alpha$-$GA_3$ antibody was detected with a secondary α-mouse IgG antibody (red), neurons are stained with a Tuj1 antibody (white) and nuclei with DAPI (blue). Insets show a magnification of the dotted white square from the merge image. Scale bars = 20 µm. **(B)** Confocal images of 8.5-week-old neural cultures treated for 21 days with 0.2 µM of $^{ch}\alpha$-$GA_1$ antibody. The arrow points to a neuron expressing $GA_{50}$-GFP and having internalized $^{ch}\alpha$-$GA_1$. Arrowheads point to round GA aggregates that do not colocalize with $^{ch}\alpha$-$GA_1$. Scale bar = 25 µm or 5 µm. **(C)** $GA_{50}$-GFP (green) and $^{ch}\alpha$-$GA_3$ antibody (red) channels of a 100x confocal image showing a cell expressing $GA_{50}$-GFP and having internalized $^{ch}\alpha$-$GA_3$ antibody. The 20 optical sections (Z-stacks) are represented in the vertical and horizontal orthogonal panels. Scale bars = 10 µm and 1 µm in inset.

**Fig. S9. Poly-GA and antibodies form large hetero-complexes in long-term treated neuronal culture**

**(A)** GFP channel of non-treated human neurons expressing $GA_{50}$-GFP at the 3, 7 and 21 days time points. Scale bar = 20 µm. **(B)** Quantification of intracellular $GA_{50}$-GFP structures normalized to the number of nuclei for samples treated with $^{ch}\alpha$-$GA_3$. 36 images per well, 3-6 wells per condition. **(C)** Quantification of $GA_{50}$-GFP extracellular structures normalized to the number of nuclei for samples treated with $^{ch}\alpha$-$GA_3$. Manual counting of 5 images per well, 3-6 wells per condition. Statistical significance tested with the beta-binomial test: relevant significance indicated as ns P >0.05, ** P ≤ 0.01, *** P ≤ 0.001. **(D)** Confocal imaging of a $GA_{50}$-GFP-expressing neural culture showing an $^{ch}\alpha$-$GA_3$ extracellular structures colocalizing with $GA_{50}$-GFP. Antibody is detected with α-mouse-Alexa 647. **(E)** Nuclei quantification, 36 images per well, 3-6 wells per condition. **(F)** Western blot of the insolubility assay supernatant fractions probed against GFP and actin as a loading control. **(G)** Representative blots of a filter retardation assay with resuspended pellet fractions. $GA_{50}$-GFP was detected with an α-GFP antibody. (**H**) Quantification of the filter retardation blot for human neurons treated during 3 or 7 days with $^{ch}\alpha$-$GA_3$ or isotype control. The intensity of each dot was normalized to the non-treated samples at 3 days. Each dot represents one replicate. Mann and Whitney test * P ≤ 0.05, ** P ≤ 0.01.

**Fig. S10. Correlative light-electron microscopy of purified $GA_{50}$-GFP aggregates**

**(A-E)** Representative correlative light-electron microscopy (CLEM) of purified aggregates from antibody-treated $GA_{50}$-GFP-expressing HEK293T cellular extracts. **(A)** Brightfield microscopy overview of the entire coverslip with gold letter grid for orientation purposes. **(B-D)** Immunofluorescence imaging of purified $GA_{50}$-GFP aggregates of the red inset from (A); **(B)** GFP (aggregate) channel; **(C)** far red (antibody) channel and **(D)** merge including brightfield. **(E)** CLEM performed on the (B-C) field of view combining immunofluorescence imaging and SEM imaging. Note the high-resolution SEM image inset of a $GA_{50}$-GFP aggregate from the upper-right quadrant of (B). **(F-H)**, **(I-K)** and **(L-N)** immunofluorescence images of purified $GA_{50}$-GFP aggregates from non-treated, α-$GA_1$- or α-$GA_3$-treated $GA_{50}$-

GFP-expressing HEK293T cellular extracts, respectively. **(F, I, L)** GFP (aggregate) channel, **(G, J, M)** far red (antibody) channel and **(H, K, N)** corresponding merges including brightfield.

**Fig. S11. Pharmacokinetics of DPR antibodies in C9$^{450}$ mice**

**(A)** Diagram of pharmacokinetic experimental design in C9$^{450}$ mice. "h" refers to hours and "d" to days. At least 3 mice were bled and taken down at each time point. **(B-D)** Plasma and brain antibody levels at different times post-injection are shown for $\alpha$-GA$_1$ **(B)**, $\alpha$-GA$_3$ **(C)**, and $\alpha$-GP$_1$ **(D)**. **(E)** C9$^{450}$ brain sections from PBS, $\alpha$-GA$_1$, $\alpha$-GA$_3$, or $\alpha$-GP$_1$ treated mice were immunostained for human DPR antibodies (Alexa Fluor® 488 AffiniPure Goat Anti-Human IgG, red) and poly-GA (Rb4334, green). Scale bar represents 5 μm.

**Fig. S12. $\alpha$-GA antibodies increase poly-GA level in insoluble fractions after ultracentrifugation**

**(A)** Schematic representation of a protocol to separate soluble and insoluble protein fractions from mouse brains. C9$^{450}$ brains treated with PBS, $^{ch}\alpha$-GA$_1$, $^{ch}\alpha$-GA$_3$, or $^{ch}\alpha$-GP$_1$ antibodies were separated into 2 % SDS (Fraction 1) or urea 7 M (Fraction 2) soluble fractions. **(B)** Total *C9ORF72* RNA levels determined by qRT-PCR in 7 month-old C9$^{450}$ mice treated either with PBS, $^{ch}\alpha$-GA$_1$, $^{ch}\alpha$-GA$_3$, or $^{ch}\alpha$-GP$_1$ antibodies. **(C-D)** Quantification of poly-GP **(C)** and poly-GA **(D)** by immunoassay in the 2 % SDS soluble Fraction 1. **(E)** Quantification of poly-GA by immunoassay in the urea 7 M fraction after ultracentrifugation (Fraction 2). **(F)** Schematic representation of a fractionation protocol after antibodies were introduced *in vitro* in mouse brain lysates. **(G-H)** Quantification of poly-GA in 2 % SDS **(G)** or urea 7 M **(H)** by immunoassay using lysates mixed with PBS, IgG isotype, $^{ch}\alpha$-GA$_1$, $^{ch}\alpha$-GA$_3$, or $^{ch}\alpha$-GP$_1$ antibodies at 3, 0.3, 0.03, 0.003 μM.

**Fig. S13. $^{ch}\alpha$-GA$_1$ antibody and poly-Ub proteins are detected in the sarkosyl-insoluble fraction of AAV$(G_4C_2)_{149}$ mouse brains**

**(A)** Sarkosyl-insoluble fractions (pellets) of brain homogenates from $^{ch}\alpha$-GA$_1$ or IgG isotype control-treated AAV$(G_4C_2)_n$ mice analyzed by filter retardation assay. Re-suspended pellets were passed through a 0.2 µm nitrocellulose membrane and probed with secondary HRP-conjugated goat α-mouse antibody only (to detect the chimeric treatment antibody). **(B)** Signal intensity quantification of the filter retardation assay from A. **(C)** Sarkosyl insoluble fractions analyzed by western blot to detect poly-ubiquitinated proteins via rabbit α-UBK48 primary antibody. **(D)** Signal intensity quantification of the poly-ubiquitin western blot from **C**. Each square on the graphs represents one mouse. Horizontal lines represent the mean ± SD, one-way ANOVA followed by Tukey's multiple comparison test. No indication $P > 0.05$, *** $P \leq 0.0001$.

**Fig. S14. Poly-GA and poly-GR aggregates in brains of AAV$(G_4C_2)_{149}$ mice treated with $^{ch}\alpha$-GA$_1$ antibody or IgG isotype control**

**(A, B)** Immunofluorescence for poly-GA (green) and poly-GR (red) **(A)**, or poly-GP (red) **(B)**, in the motor cortex of 4-month-old AAV-$(G_4C_2)_2$ or AAV-$(G_4C_2)_{149}$ mice treated with $^{ch}\alpha$-GA$_1$ or isotype control. Scale bar: 25 µm. **(C-F)** Quantifications of percent area occupied by poly-GA aggregates detected with a rabbit anti-poly-GA antibody in cortex **(C, E)** and hippocampus **(D, F)** of 4 month-old **(C-D)** and 10 month-old **(E-F)** AAV-$(G_4C_2)_2$ or AAV-$(G_4C_2)_{149}$ mice treated with $^{ch}\alpha$-GA$_1$ or IgG isotype control. **(G, H)** Quantification of poly-GR levels by immunoassay **(G)** and immunohistochemistry **(H)** in cortex of 4-month-old AAV-$(G_4C_2)_2$ or AAV-$(G_4C_2)_{149}$ mice treated with $^{ch}\alpha$-GA$_1$ or IgG isotype control. Error bars represent the mean ± SD, one-way ANOVA followed by Tukey's multiple comparison test. Not significant (ns).

**Fig. S15. pTDP-43 aggregates in 12-month old AAV($G_4C_2$) mice treated with $^{ch}\alpha$-GA$_1$ antibody or IgG isotype control**

**(A)** Immunohistochemistry staining for phospho-TDP-43 (pTDP-43) in the cortex of 12-month-old AAV($G_4C_2$) mice after 10 months of treatment with $^{ch}\alpha$-GA$_1$ or IgG isotype control. **(B)** Quantification of the number of pTDP-43 inclusions in cortex of 12-month-old AAV($G_4C_2$) mice. Error bars represent the mean ± SD, one-way ANOVA followed by Tukey's multiple comparison test. Not significant (ns).

**Fig. S16. Chronic administration of human-derived DPR antibodies is well tolerated in two C9ORF72 mouse models**

**(A-C)** Serum antibody titers of $^{ch}\alpha$-GA$_1$ **(A)**, $^{ch}\alpha$-GA$_3$ **(B)**, or $^{ch}\alpha$-GP$_1$ **(C)** quantified by ELISA every 8 weeks. **(D, E)** Mean body weights over 16 months for female **(D)** and male **(E)** $C9^{450}$ mice treated with $^{ch}\alpha$-GA$_1$, $^{ch}\alpha$-GA$_3$, $^{ch}\alpha$-GP$_1$ or PBS. Error bars at 17 ,36, 58 and 71 weeks represent standard deviation, one-way ANOVA followed by Tukey's multiple comparison test, ns = not significant. (**F**) Distance traveled in the open-field test by 13-month-old males treated with PBS, $^{ch}\alpha$-GA$_1$, $^{ch}\alpha$-GA$_3$ and $^{ch}\alpha$-GP$_1$ antibodies ($n \geq 10$ per group). Bars represent the mean ± SD, Kruskal-Wallis test followed by Dunnett's multiple comparison tests. Not significant (ns) $P > 0.05$, ** $P \leq 0.01$ and *** $P \leq 0.001$. **(G, H)** Mean body weights over 16 months for female **(G)** and male **(H)** $(G_4C_2)_{149}$ and $(G_4C_2)_2$ mice treated with $^{ch}\alpha$-GA$_1$ or IgG isotype control. Error bars at 9, 19, 29 and 39 weeks represent standard deviation, one-way ANOVA followed by Tukey's multiple comparison test, ns = not significant.

| Human Antibody | EC50 (nM) | | | | | Kinetics for main target (*) | | | IHC human | IHC mouse | Immunofluorescence: NSC-34 cells | | | | |
|---|---|---|---|---|---|---|---|---|---|---|---|---|---|---|---|
| | $(GA)_{15}$ | $(GP)_{15}$ | $(GR)_{15}$ | $(PR)_{15}$ | $(PA)_{15}$ | $K_D \pm$ SD (M) | $k_a \pm$ SD ($M^{-1}$ $s^{-1}$) | $k_d \pm$ SD ($s^{-1}$) | | | $(GA)_{50}$ | $(GP)_{47}$ | $(GR)_{50}$ | $(PR)_{50}$ | $(PA)_{50}$ |
| α-$GA_1$ | 0.2* | - | - | - | - | $(2.2 \pm 0.3) *10^{-9}$ | $(4.4 \pm 0.7) *10^{5}$ | $(9.3 \pm 0.5) *10^{-4}$ | + | + | + | - | - | - | - |
| α-$GA_2$ | 0.3* | - | - | - | - | $(1.8 \pm 0.5) *10^{-10}$ | $(1.6 \pm 0.1) *10^{5}$ | $(2.8 \pm 0.5)\ 10^{-5}$ | + | + | + | - | - | - | - |
| α-$GA_3$ | 0.3* | - | - | - | - | $(8 \pm 9) *10^{-11}$ | $(1.5 \pm 0.2) *10^{5}$ | $(1 \pm 1) *10^{-5}$ | + | + | + | - | - | - | - |
| α-$GA_4$ | 0.3* | - | - | - | - | $(3 \pm 2) *10^{-11}$ | $(2.1 \pm 0.3) *10^{5}$ | $(6 \pm 6)\ 10^{-6}$ | + | ND | + | - | - | - | - |
| α-$GP_1$ | 8.0 | 0.3* | - | - | - | $(3.6 \pm 0.2) *10^{-9}$ | $(2.5 \pm 0.2) *10^{5}$ | $(9.1 \pm 0.3) *10^{-4}$ | + | + | + | + | - | - | - |
| α-$GR_1$ | 69.1 | - | 0.1* | >200 | - | $(8.5 \pm 0.2) *10^{-10}$ | $(6.1 \pm 0.8) *10^{5}$ | $(5.2 \pm 0.6) *10^{-4}$ | - | - | - | - | - | - | - |
| α-$PR_1$ | 6.3 | - | 22 | 3.8* | - | $(1.7 \pm 0.1) *10^{-9}$ | $(1.7 \pm 0.1) *10^{5}$ | $(3.0 \pm 0.5) *10^{-4}$ | - | ND | - | - | - | +/- | - |
| α-$PR_2$ | - | - | - | 12.8* | - | $(1.1 \pm 0.2) *10^{-9}$ | $(2.0 \pm 0.3) *10^{6}$ | $(2.0 \pm 0.2) *10^{-3}$ | - | ND | - | - | - | + | - |
| α-$PR_3$ | 55.1 | - | - | 3.5* | - | $(7.0 \pm 0.2) *10^{-10}$ | $(5.9 \pm 0.1) *10^{5}$ | $(4.11 \pm 0.04) *10^{-4}$ | ND | ND | - | - | - | + | - |
| α-$PA_1$ | > 100 | - | - | - | 0.1* | $(2.3 \pm 0.1) *10^{-9}$ | $(6.9 \pm 0.3) *10^{5}$ | $(1.60 \pm 0.02) *10^{-3}$ | - | - | - | - | - | - | + |
| α-$PA_2$ | 10.4 | - | - | - | 0.4* | $(2.21 \pm 0.05) *10^{-9}$ | $(7.7 \pm 0.9) *10^{5}$ | $(1.7 \pm 0.2) *10^{-3}$ | - | - | +/- | - | - | - | + |

**Table S1. Characterization of human-derived antibodies against DPRs**

Affinity, kinetics and specificity of human-derived monoclonal α-DPR antibodies determined by ELISA, biolayer interferometry, immunohistochemistry (IHC) and immunofluorescence (IF). Kinetic data is shown for the primary DPR target as determined by ELISA and indicated with an asterisk (*). $K_D$ equilibrium dissociation constant, $k_a$ association rate constant and $k_d$ dissociation rate constant.

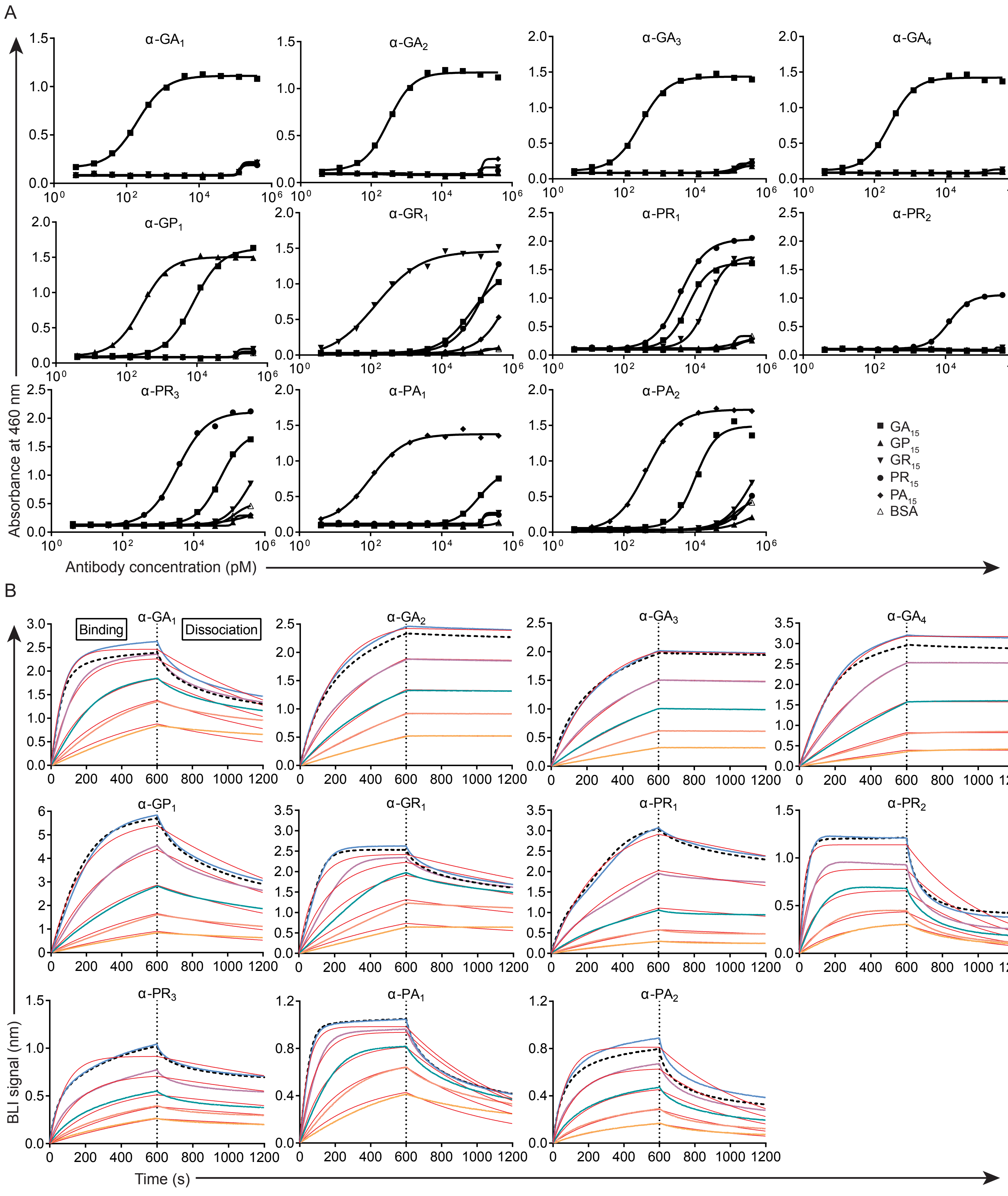


Figure S1. Specificity and binding kinetics of human-derived antibodies against DPRs

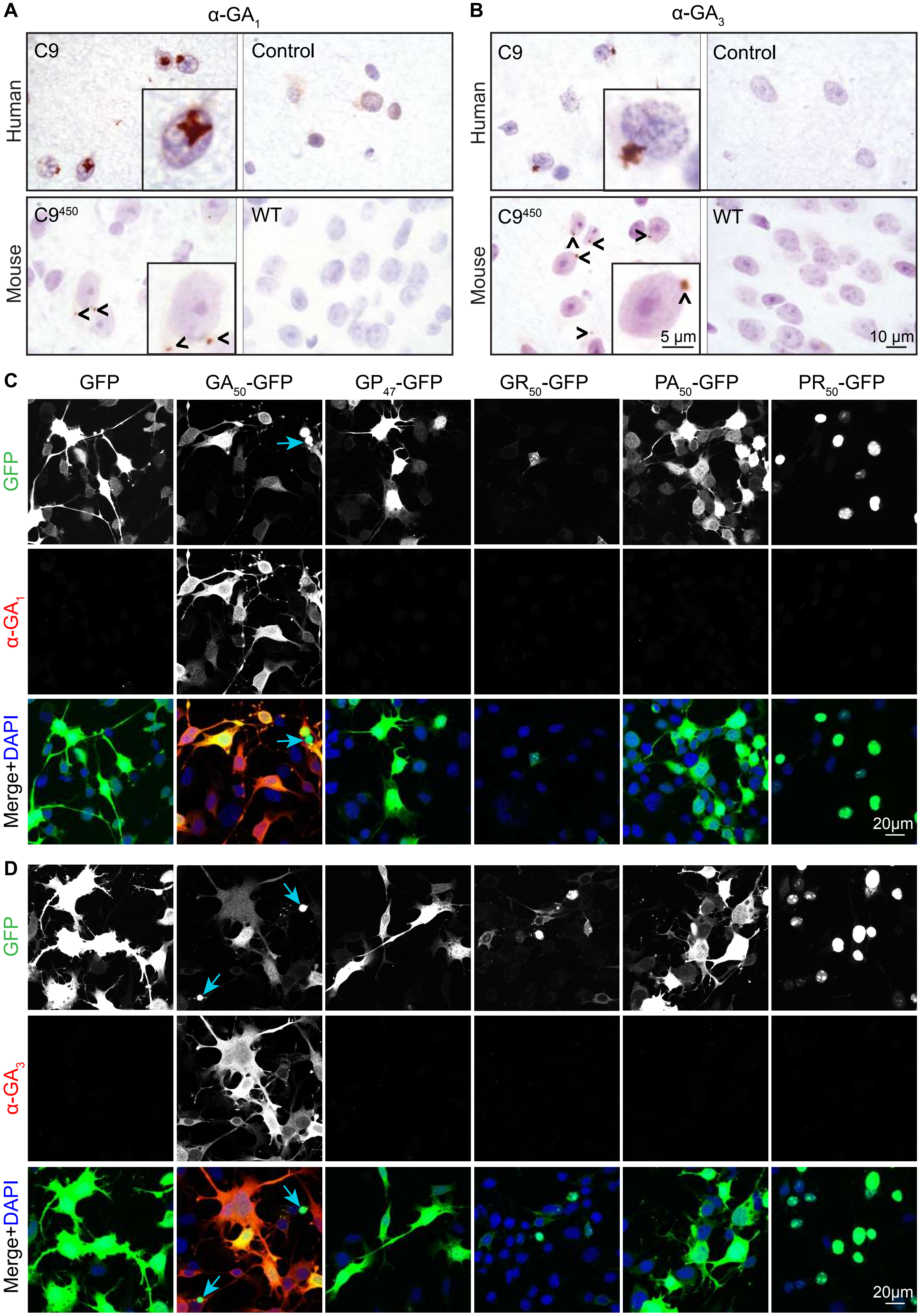


Figure S2. Characterization of α-$GA_1$ and α-$GA_3$ antibodies binding specificity in tissues and motor-neuron-like cells

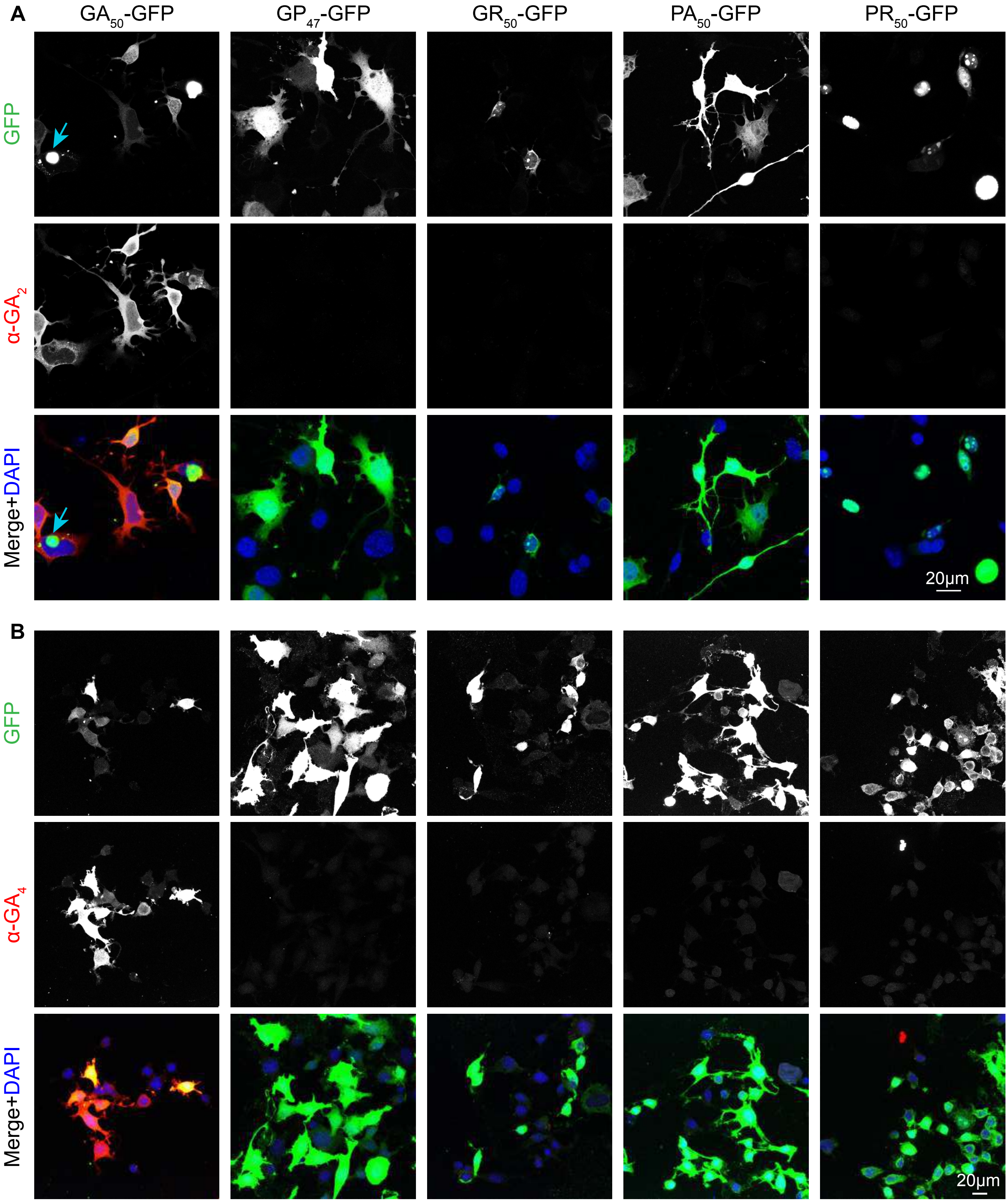


Figure S3. Characterization of α-GA binding specificity in motor-neuron like cells

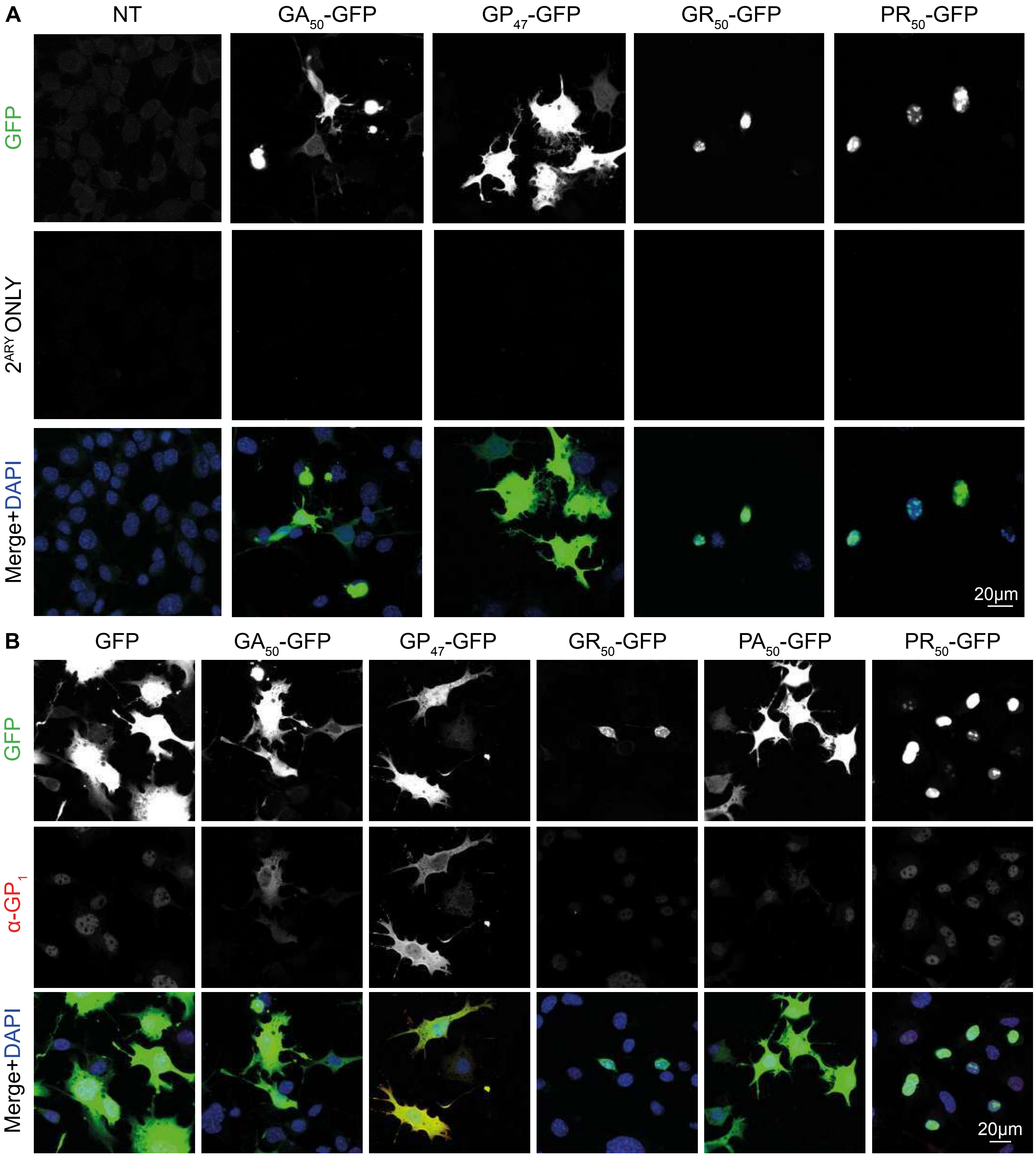


Figure S4. Characterization of α-GP binding specificity in motor-neuron like cells

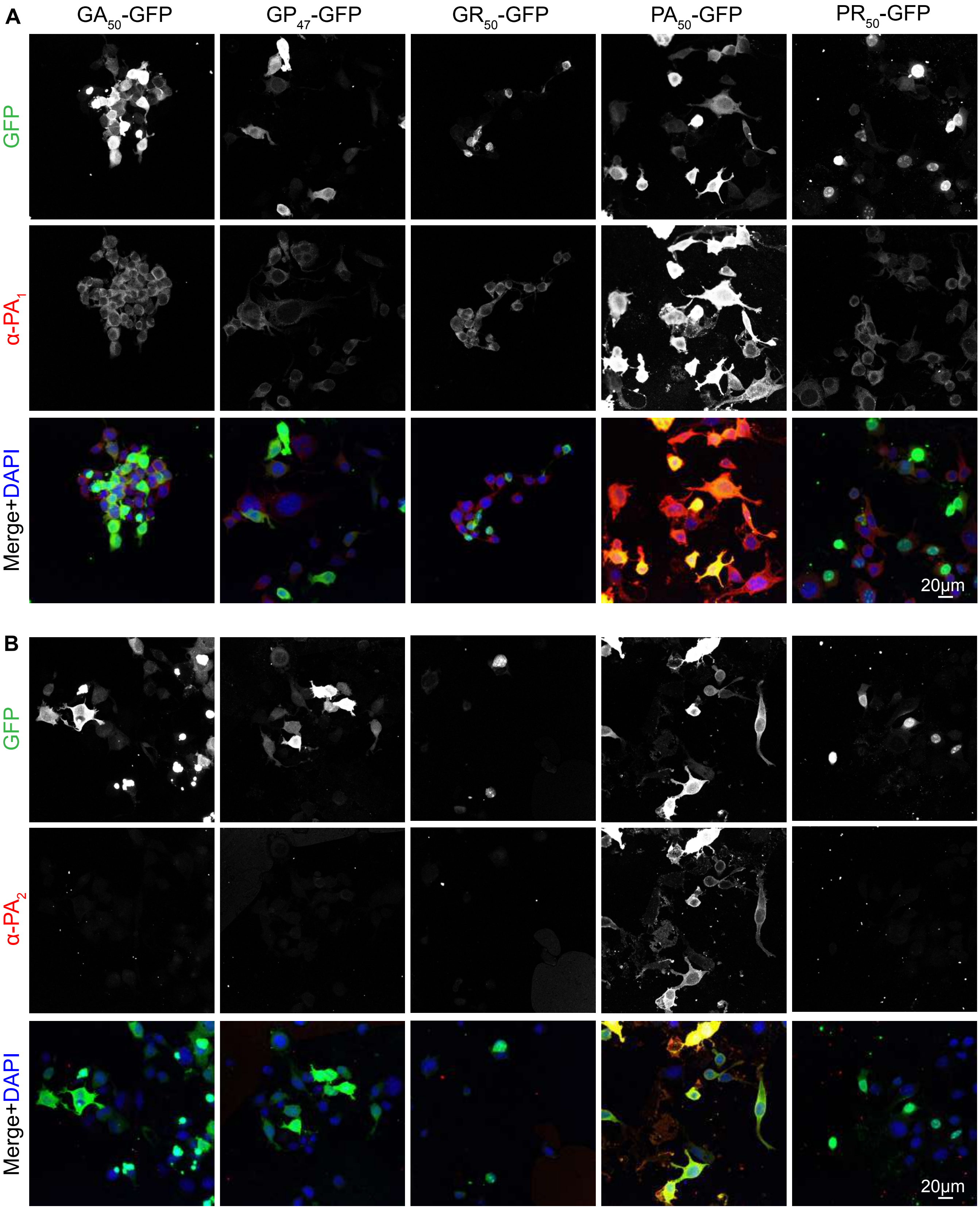


Figure S5. Characterization of α-PA binding specificity in motor-neuron like cells

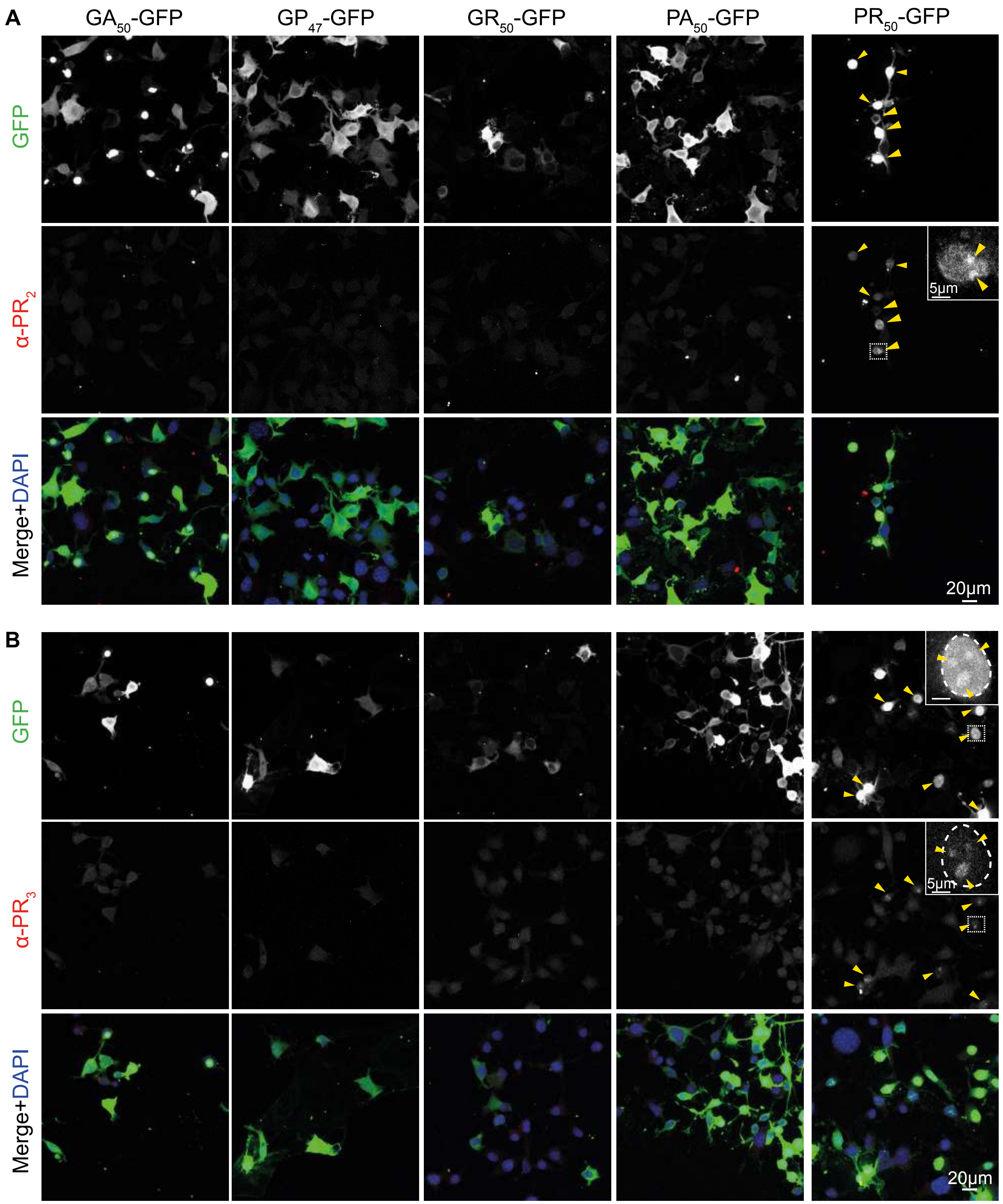


Figure S6. Characterization of α-PR binding specificity in motor neuron-like cells

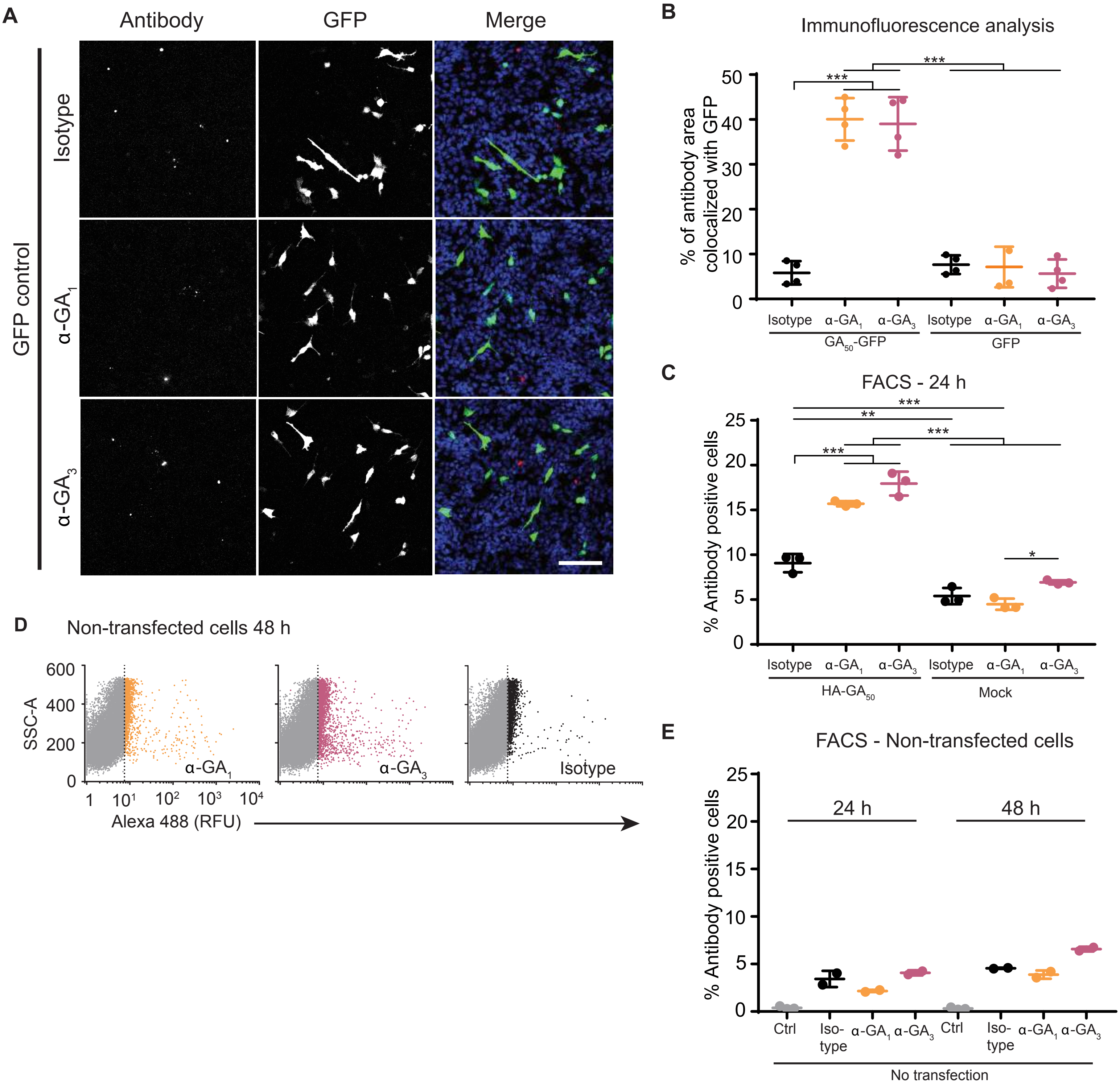


Figure S7. Intracellular expression of $GA_{50}$ increases uptake/retention of poly-GA specific antibodies in SH-SY5Y cells

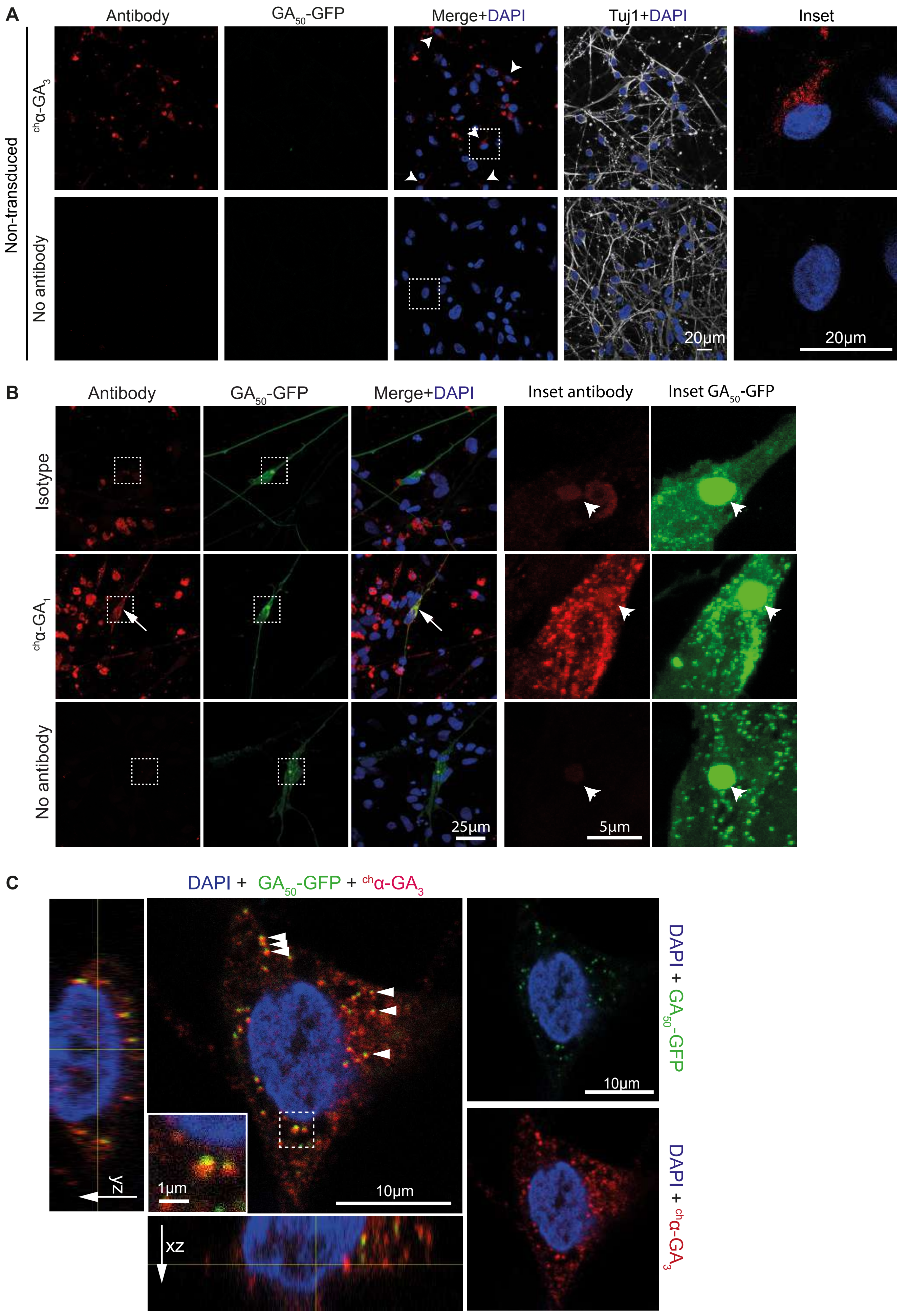


Figure S8. Antibody internalization and colocalization with $GA_{50}$-EGFP in human neural culture

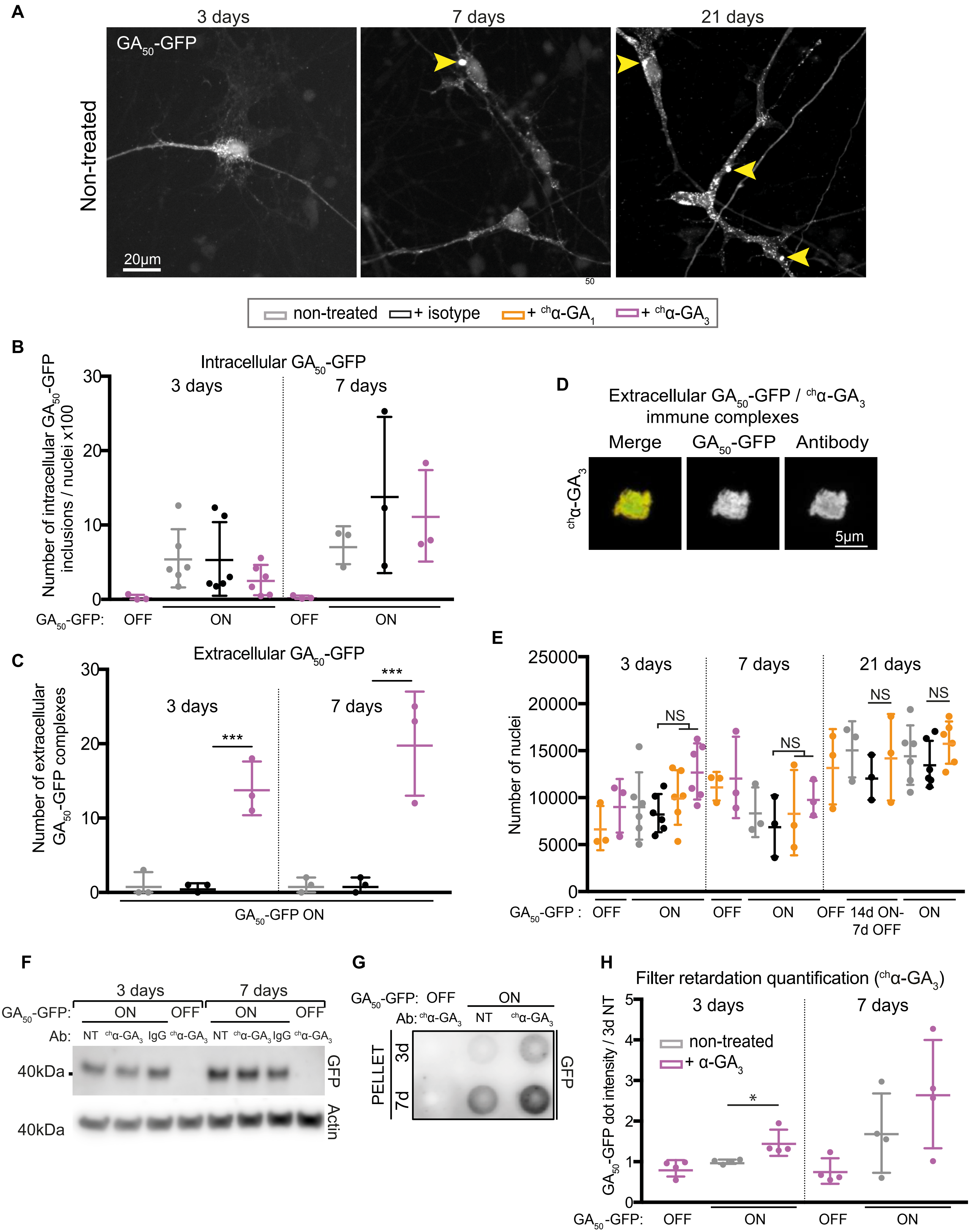


Figure S9. Poly-GA and antibodies form large hetero-complexes in long-term treated neural culture

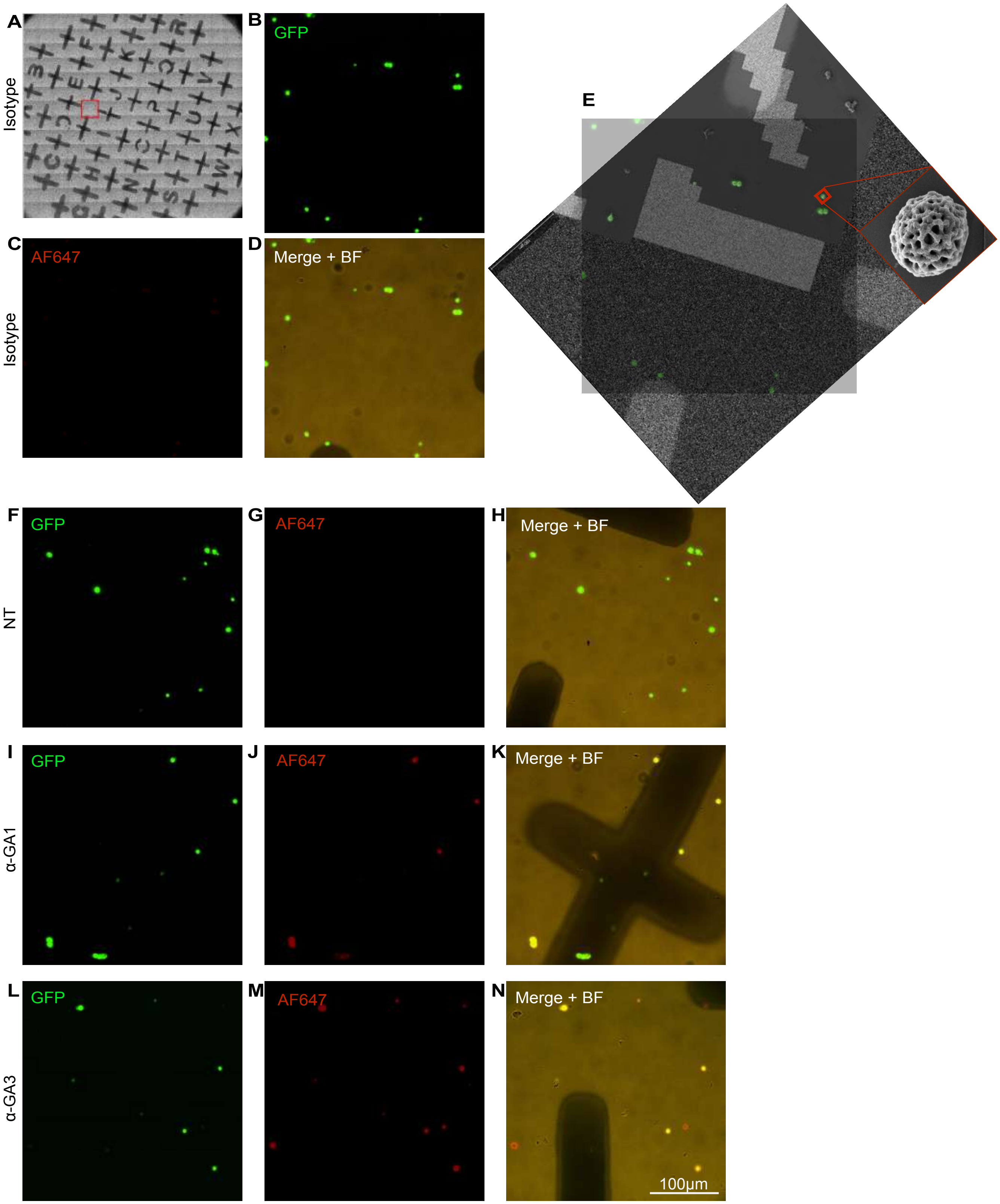


Figure S10. Correlative light-electron microscopy (CLEM) of purified $GA_{50}$-GFP aggregates

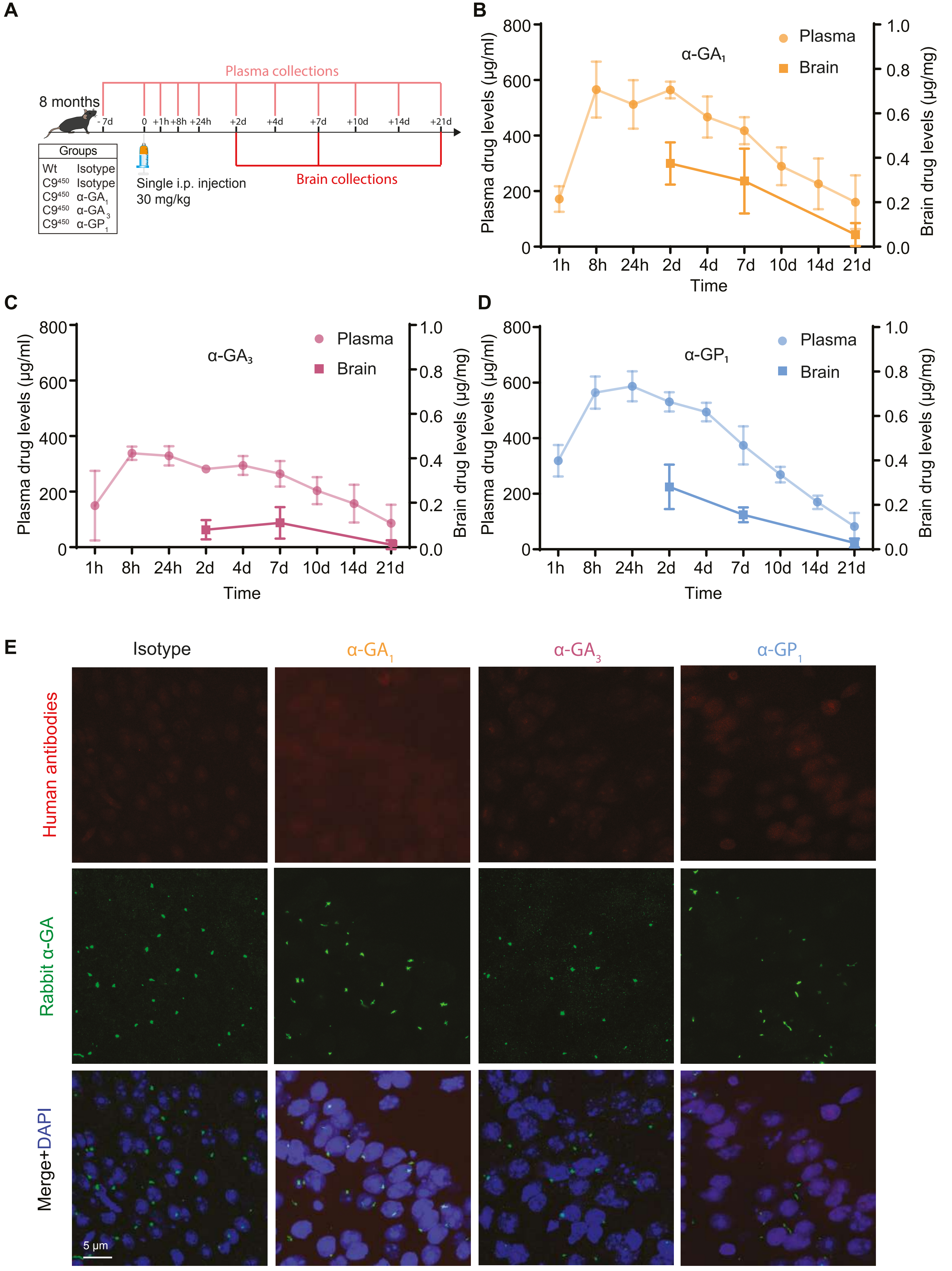


Figure S11. Pharmacokinetics and target engagement of DPR antibodies in C9[450] mice

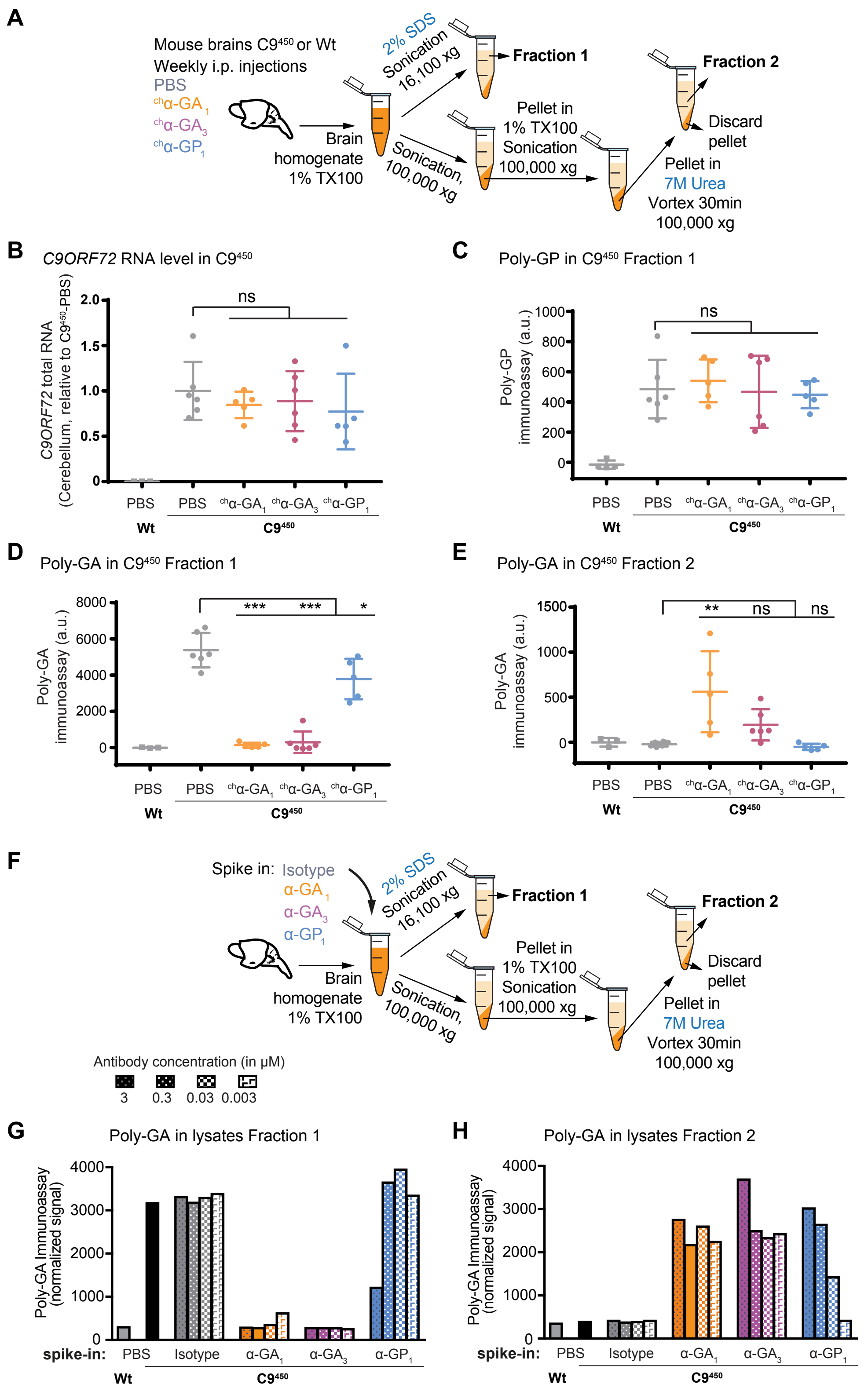


Figure S12. α-GA antibodies increase poly-GA insolubility

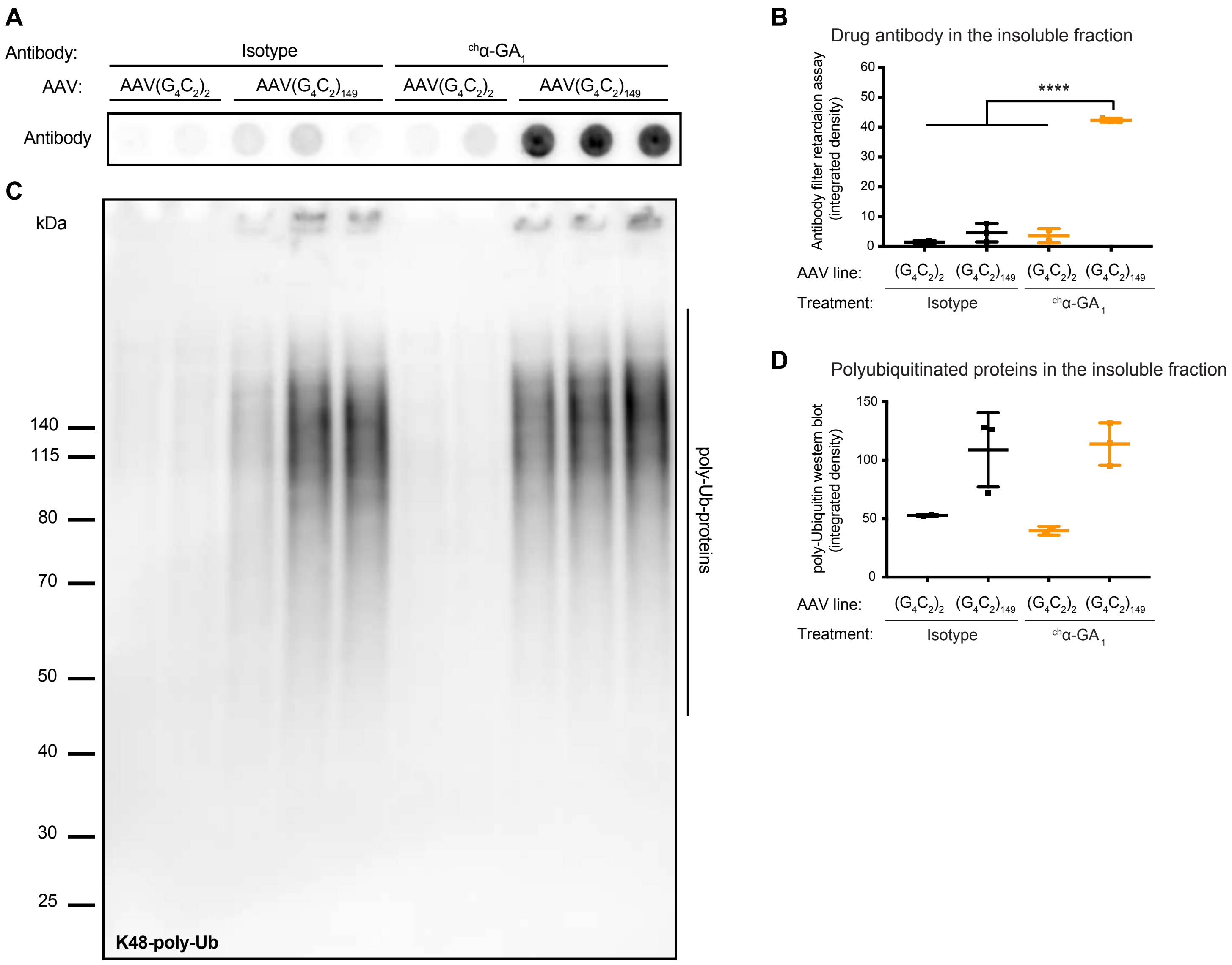


Figure S13. $^{ch}$α-$GA_1$ antibody and poly-Ub proteins are detected in the sarkosyl-insoluble fraction of $AAV(G_4C_2)_{149}$ mouse brains

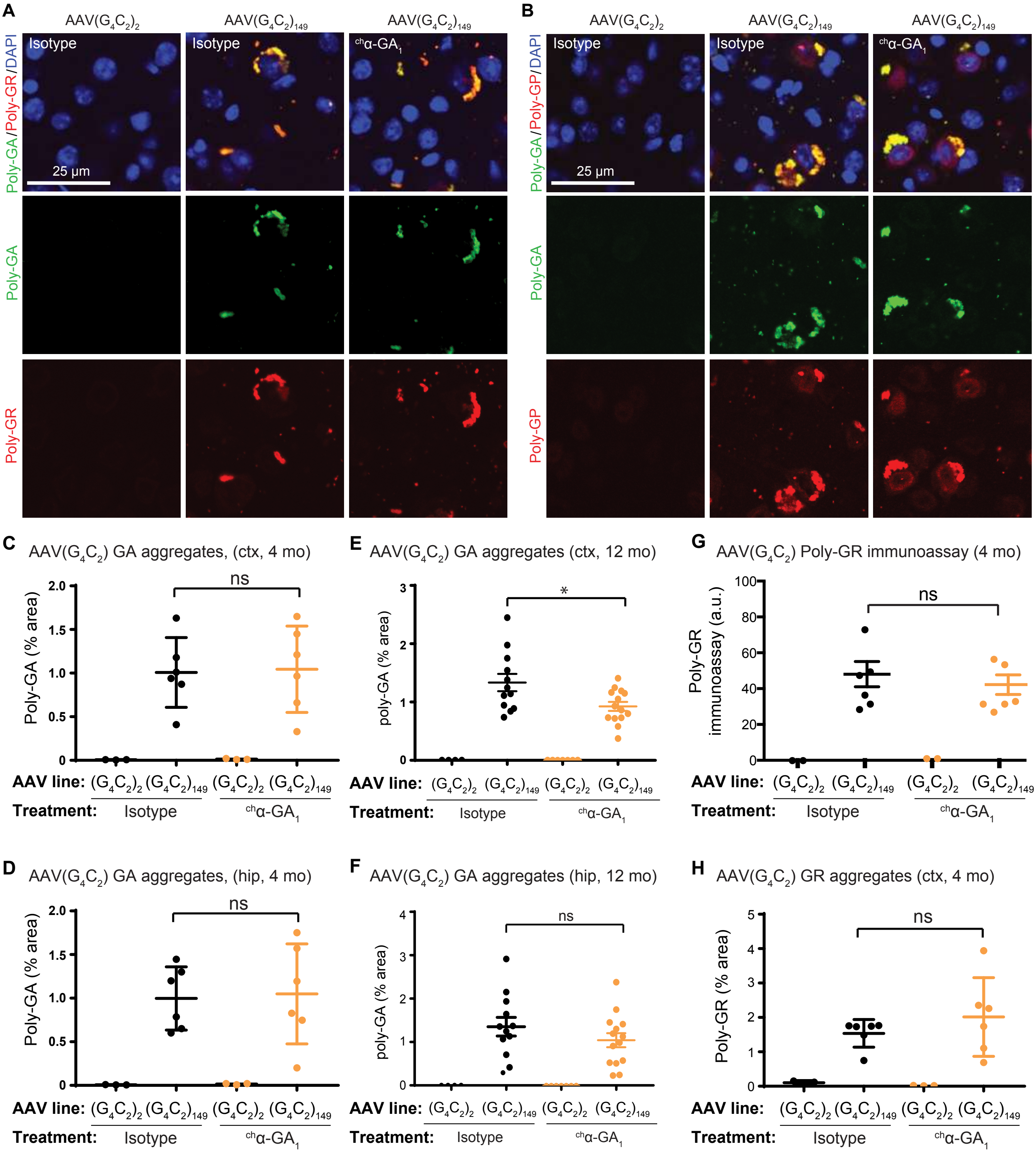


Figure S14. Poly-GA and poly-GR aggregates in brains of AAV($G_4C_2$)$_{149}$ mice treated with $^{ch}$α-GA$_1$ antibody or IgG isotype control

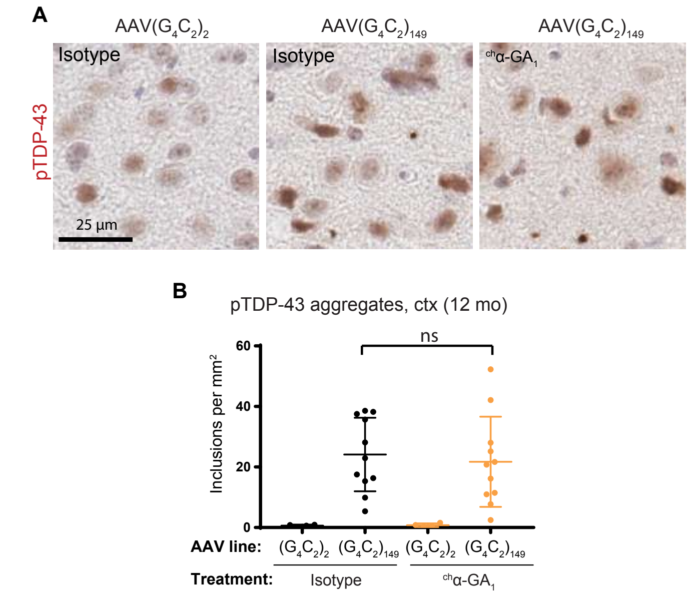


Figure S15. pTDP-43 aggregates in 12 month-old AAV($G_4C_2$) mice treated with $^{ch}\alpha$-$GA_1$ antibody or IgG isotype control

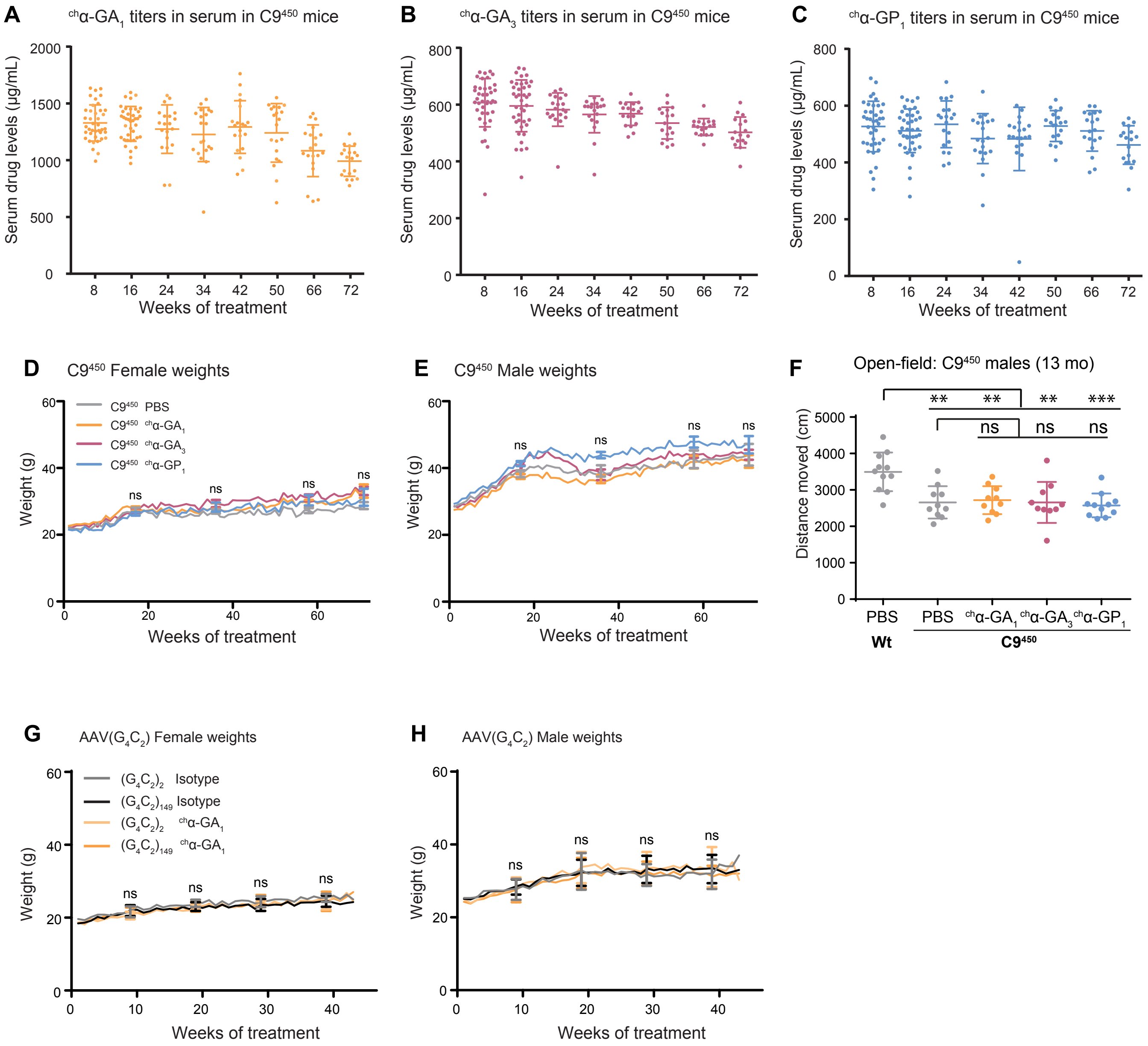


Figure S16. Chronic administration of human-derived DPR antibodies is well tolerated in two C9ORF72 mouse models
